# Large circulation of a novel vesiculovirus in bats in the Mediterranean region

**DOI:** 10.1101/2024.04.24.590417

**Authors:** Dong-Sheng Luo, Markéta Harazim, Corinne Maufrais, Simon Bonas, Natalia Martinkova, Aude Lalis, Emmanuel Nakouné, Edgard Valéry Adjogoua, Mory Douno, Blaise Kadjo, Marc López-Roig, Jiri Pikula, Zheng-Li Shi, Hervé Bourhy, Jordi Serra-Cobo, Laurent Dacheux

**Author notes:** Correspondence: Dongsheng Luo Laurent Dacheux. Present address: London School of Hygiene & Tropical Medicine, WC1E 7HT, London, UK. Present address: Institut Pasteur, Université Paris Cité, Unit Environnement and Infectious Risks, Institut Pasteur, Paris, France (Laurent Dacheux).

## Abstract

Bats are the natural reservoirs of a variety of emerging or re-emerging viruses. Among them, rabies virus (genus *Lyssavirus*, family *Rhabdoviridae*) is of the first and most iconic described in these animals. Since its first description, various new bat lyssaviruses have been regularly described. Apart from lyssaviruses, other bat rhabdoviruses have been also identified, including members of the *Vesiculovirus*, *Ledantevirus* and more recently *Alphanemrhavirus* and *Tupavirus* genera. However, the family *Rhabdoviridae* is one of the most abundant and diverse viral families, with 318 officially recognized species divided into 3 subfamilies and 46 different genera. Thus, the number of bat- associated rhabdoviruses is probably higher.

In this study, we first developed and validated a combined nested RT-qPCR technique (pan-rhabdo RT-nqPCR) dedicated to the broad detection of animal rhabdovirus. After validation, this technique was used for a large retrospective screening of archival bat samples (*n* = 1962), including blood (*n* = 816), brain (*n* = 723) and saliva (*n* = 423). These samples were collected from various bat species over a period of 12 years (2007-2019) in 9 different countries in Europe and Africa. A total of 23 samples (1.2%) from *Miniopterus schreibersii*, *Rhinolophus euryale* and *Rhinolophus ferrumequinum* bat species was found positive for rhabdovirus infection, including 17 (2.1%) blood and 6 (1.4%) saliva samples, all collected from bats originated from the Mediterranean region.

The complete virus genome sequences were obtained by next-generation sequencing for most of the positive samples. Molecular and phylogenetic analysis of these sequences demonstrated that these virus isolates, named Mediterranean bat virus (MBV), were closely related, and represented a new species *Vesiculovirus mediterranean* within the *Vesiculovirus* genus. MBV was more specifically related to the other bat vesiculoviruses previously described in China and North America, together clustering into a distinct group of bat viruses within this genus. Interestingly, our results suggest that MBV is widely distributed, at least in the West part of the Mediterranean region, where it can act as an arbovirus infecting and circulating in multiple bat species. These findings expand the host range and the viral diversity of bat vesiculoviruses and pave the way for further investigations to determine the route of transmission and the dynamic of diffusion of these viruses into bat colonies, as well as to evaluate their potential hazard for public health.

## 1 Introduction

Bats belong to the order Chiroptera, which represents the second largest mammalian order, with 1466 extant species so far, belonging to 21 bat families (https://www.mammaldiversity.org/). These animals have many specific biological and ecological characteristics, including various unique living habits and extensive geographical distribution (Teeling et al., 2005; Jebb et al., 2020; Zhai et al., 2020). In addition, bats are attracting growing interest from the scientific community, particularly in terms of public health. Indeed, in the past few decades, hundreds of virus species belonging to various families have been detected or isolated in bats, suggesting that these animals represent a major animal reservoir (Calisher et al., 2006; Hayman, 2016; Van Brussel and Holmes, 2022; Weinberg and Yovel, 2022). Among them, numerous emerging or re-emerging infectious diseases were demonstrated or suspected to be connected with bats, including coronavirus-based diseases with severe acute respiratory syndrome (SARS) (Li, 2005; Ge et al., 2013), Middle-East respiratory syndrome (MERS) (Chan et al., 2015) and more recently the coronavirus disease 2019 (COVID-19) pandemic (Zhou et al., 2020), as well as fatal hemorrhagic diseases caused by filoviruses (Ebola and Marburg viruses) (Leroy et al., 2005; Towner et al., 2009) and henipaviruses (Hendra and Nipah viruses) (Ang, 2018; Kessler et al., 2018; Mbu’u et al., 2019). Furthermore, as the research on bat-associated viruses advances, this viral diversity continues to expand, particularly in other families of viruses (Van Brussel and Holmes, 2022). However, some of these virus families still remain poorly investigated such as the *Rhabdoviridae* family.

This virus family belongs to the order *Mononegavirales* and encompasses single-stranded negative RNA viruses. According to the last update of the International Committee of Taxonomy of Viruses (ICTV), this family encompasses 318 different species distributed among 3 subfamilies and 46 genera (Walker et al., 2022b; Kuhn et al,, 2022). They are typically enveloped virions with bullet-shaped or bacilliform morphology, and are reported to be in the range of 100–460 nm in length and 45–100 nm in diameter. The virion is generally characterized by a linear single stranded negative-sense RNA genome of approximately 10–16 kb with five canonical genes encoding the nucleoprotein (N), the phosphoprotein (P), the matrix protein (M), the glycoprotein (G), and the RNA-dependent RNA polymerase (L) (Walker et al., 2022b). These viruses exhibit a large ecological diversity (Walker et al., 2015), with members able to infect plants (Dietzgen et al., 2020) but also animals such as birds (Luo et al., 2021b), reptiles (Vasilakis et al., 2019) or fishes (Walker et al., 2022a). Remarkably, rhabdoviruses were often isolated from insects or are categorized as arboviruses (Vasilakis et al., 2014; Contreras et al., 2017; Walker et al., 2022b).

Mammals represent also important animal reservoirs or susceptible hosts of rhabdoviruses (Kuzmin et al., 2009; Marston et al., 2018; Walker et al., 2022a, 2022b). Among them, bats have been associated so far with four different rhabdovirus genera. The first and the most emblematic one is the genus *Lyssavirus*, which includes rabies virus (RABV) (Rupprecht et al., 2017; Marston et al., 2018). Indeed, RABV was the first described bat virus, and nearly all the different lyssavirus species or tentative species have been isolated from bats (Fooks et al., 2017), which suggests that these animals are their original and natural reservoir (Badrane and Tordo, 2001). The two other main genera are *Vesiculovirus* and *Ledantevirus*, with bat viruses reported in North America and China (Ng et al., 2013; Xu et al., 2018; Luo et al., 2021a) or in several countries of Africa, Asia, Europe or North America (Calisher et al., 2006; Ghedin et al., 2013; Kading et al., 2013; Binger et al., 2015; Blasdell et al., 2015; Goldberg et al., 2017; Lelli et al., 2018; Li et al., 2022), respectively. Interestingly, many ledanteviruses have been also isolated from arthropods feeding on bats, suggesting that they can act as arboviruses (Goldberg et al., 2017; Bennett et al., 2020). Recently, one novel rhabdovirus, Sodak rhabdovirus 2 (SDRV2), was identified in North American bats, and was classified as a member of the *Alphanemrhavirus* genus (Hause et al., 2020). In addition, an individual virome analysis of Chinese bats identified putative new virus members among the *Tupavirus* genus (Wang et al., 2023). All together, these data suggest that bats are playing an important role in rhabdovirus diffusion and maintaining, with potential risks of human exposure of bat rhabdoviruses (Binger et al., 2015; Li et al., 2022), and that the rhabdoviruses diversity in these animals remains to be explored.

Due to the large genetic diversity within the family *Rhabdoviridae*, the development of generic molecular tools for rhabdovirus screening, with a satisfactory balance between specificity and sensitivity, remains a key challenge. Some innovative approaches have been conducted (Dacheux et al., 2010), but most of existing tools rely on PCR techniques, dedicated to a specific species or genera such as lyssaviruses (Dacheux et al., 2016; Luo et al., 2021a), or with an extended spectrum of detection (Aznar-Lopez et al., 2013; Conrardy et al., 2014; Fischer et al., 2014). Despite these efforts, the spectrum of detection proposed by these techniques remains limited and needs further improvement.

To achieve this goal, our study aimed to develop a combined nested RT-qPCR technique (pan-rhabdo RT-nqPCR) dedicated to the broad detection of animal rhabdoviruses by targeting a conserved region of the polymerase (L) gene. This technique was validated on an extensive list of diverse rhabdoviruses and compared to the other previous published methods, showing a significant gain in terms of specificity and sensitivity. After validation, we applied the pan-rhabdo RT-nqPCR to screen a large collection of archive bat samples collected from various species in different countries in Europe and Africa. Based on this screening, we were able to identify and characterize a new species among the genus *Vesiculovirus*, recognized by ICTV under the name of *Vesiculovirus mediterranean*, which circulate in three bat species originated from the Mediterranean region These results contribute to expand our understanding on bat rhabdovirus, but also provided a useful tool for rhabdovirus screening in bats and other animal reservoirs.

## 2 Materials and Methods

### 2.1 Ethics approval statement

Bat samples analyzed in this study were part of biocollections previously constituted from Africa (Algeria, Central African Republic, Egypt, Guinea, Ivory Coast and Morocco) and Europe (Czechia, France and Spain) in the framework of other studies. Handling and sampling protocols of individuals were covered by specific authorities of the related countries. Sample collections from Guinea and Ivory Coast were approved by the Health Ministry (2013/PFHG/05/GUI) and the Société de Développement des Forêts – SODEFOR (N°00991-16), respectively. Both were also approved by Comité Cuvier - Animal Ethics Committee of the French National Museum of Natural History (MNHN) (N°68-0009). No specific authorization was required in Central African Republic, regarding the non-protective status of bats in this country. Bat samples from Algeria, Egypt and Morocco were authorized by the respective Forest Ministries. Sampling of bats in Czechia were approved by the Agency for Nature Conservation and Landscape Protection of Czechia and complied with Czech Law No. 114/1992 on Nature and Landscape Protection. Bat sample collection in Spain was authorized by the Spanish Regional Committee for Scientific Capture from Balearic Islands and Catalonia, and by the Ethical Committee of the University of Barcelona. Lastly, samples collection from France was performed in the framework of the mission of surveillance of rabies performed by the National Reference Center for Rabies at Institut Pasteur, Paris.

### 2.2 Samples collection

A large retrospective bat sample collection was used for this study. This sample collection included a total of 1962 samples encompassing 423 saliva swabs (21.6%), 816 blood pellet samples (41.6%) and 723 brains biopsies (36.8%), collected in 9 different countries in Europe (Czechia, France and Spain) and Africa (Algeria, Central African Republic, Egypt, Guinea and Morocco) from 2007 to 2019 (Tables 1 and S1, Figure 1). These bat samples belonged to 51 species of 24 genera among eight different families, with the frugivorous bat family Pteropodidae, and the insectivorous bat families Emballonuridae, Hipposideridae, Miniopteridae, Molossidae, Rhinolophidae, Rhinopomatidae and Vespertilionidae (Tables 2 and S1). For all samples originating outside Fance, shipment was done to Institut Pasteur, Paris, in dry ice, in RNAlater (ThermoFisher) and/or in TRIzol (ThermoFisher). Upon reception, all samples were stored in -80℃.

**Figure 1.**
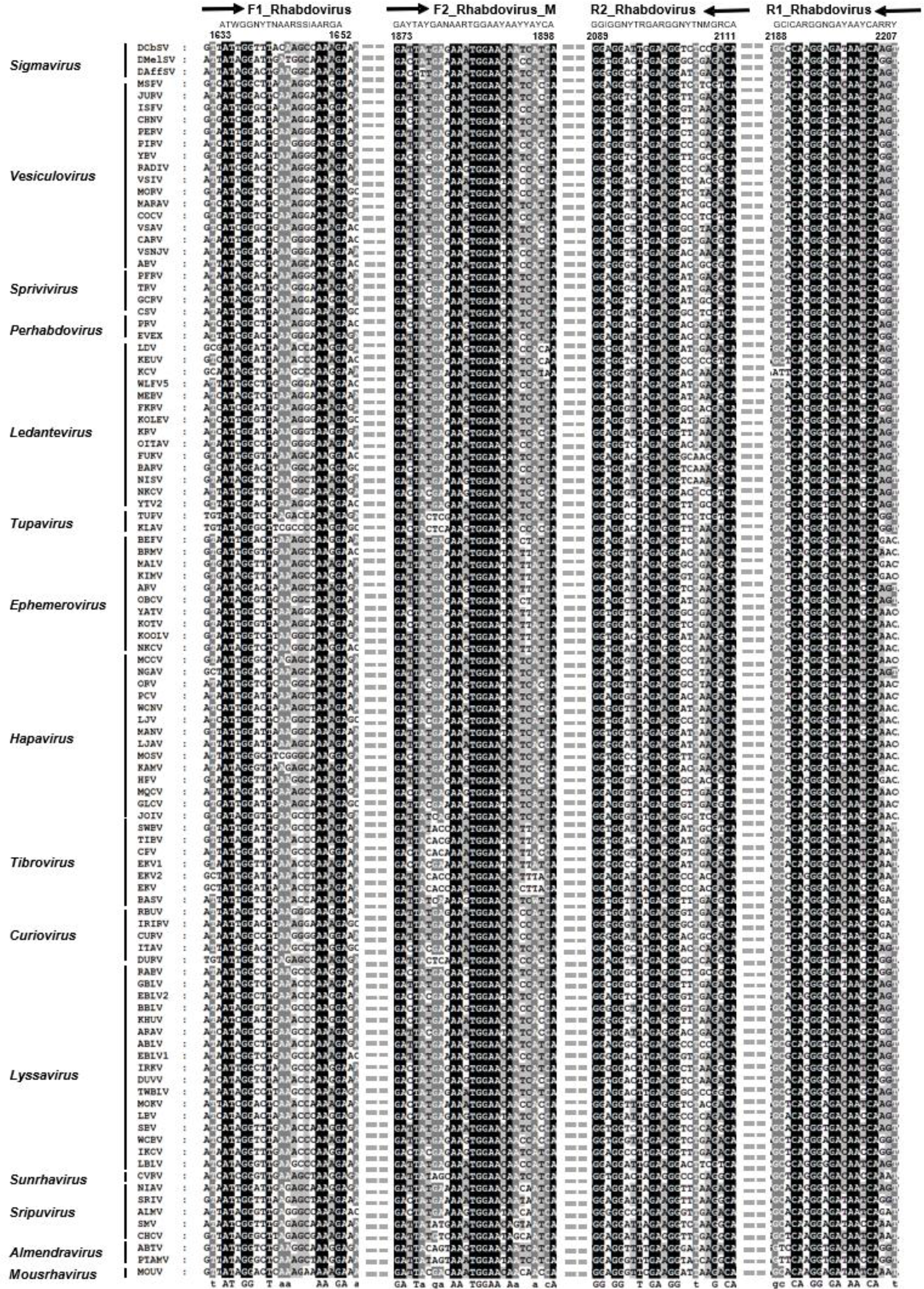
Multiple alignment of 103 amino-acid polymerase sequences of rhabdoviruses belonging to 14 different genera within the family *Rhabdoviridae*, with the position and the sequences of the four primers F1_Rhabdovirus, R1_Rhabdovirus, F2_Rhabdovirus-M and R2_Rhabdovirus selected for the pan-rhabdo RT-nqPCR. The first PCR round is based on the primers F1_Rhabdovirus (forward, 5’- ATWGGNYTNAARSSIAARGA-3’) and R1_Rhabdovirus (reverse, 5’- RYYTGRTTRTCNCCYTGIGC-3’), whereas the nested PCR round is based on primer F2_Rhabdovirus-M (forward, 5’-GAYTAYGANAARTGGAAYAAYYAYCA-3’) and R2_Rhabdovirus (reverse, 5’-TGYCKNARNCCYTCYARNCCICC-3’). The nucleotide code applied is : R=A/G, Y=C/T, M=A/C, K=G/T, S=G/C, W=A/T, H=A/T/C, B=G/T/C, V=G/A/C, D=G/A/T, N=A/T/C/G and I (Hypoxanthine). The oligonucleotide sequence of each primer is indicated in bold, together with its name and an arrow indicating the sense direction. Sequence identity of the primers based on the multiple alignment is highlighted in black. The dot lines between the different blocks of sequence represents omitted regions. The position of the primers is indicated according to the nucleotide sequence of polymerase gene of Drosophila obscura sigmavirus (DObSV) (GenBank accession number NC022580). Virus acronyms and genera are indicated in the left of the figure. The multiple alignment was performed with ClustalW, version 2.0.

Other field or laboratory samples, as well as virus isolates used in this study for the validation method step were obtained from the biocollection archives housed in the National Reference Center for Rabies and in the National Reference Center for Arboviruses (former), both at Institut Pasteur Paris, France.

### 2.3 RNA extraction and cDNA synthesis

RNA extraction from saliva samples stored in RNAlater was performed with High Pure Viral RNA Kit (Roche), both according to the manufacturer’s instructions, using 200 µL for each sample (Luo et al., 2021a). Brain and blood pellet samples were extracted with TRIzol using Direct-zol RNA MiniPrep (Zymo Research) according to the manufacturer’s instructions. Purified RNA was stored at -80°C before use. The synthesis of cDNA was conducted with Superscript III reverse transcriptase kit (Invitrogen), following the manufacturer’s instructions. For each sample, a total of 8 µL RNA was used (Pallandre et al., 2020; Luo et al., 2021a).

### 2.4 Primer designs

The design of the primers for pan-rhabdovirus screening was performed on a dataset of 103 polymerase sequences from animal rhabdoviruses belonging to 14 genera within the *Rhabdoviridae* family available in GenBank (Figure 2) (Table S2). Criteria of selection of the sequences were based on animal host (insect, fish, reptile, bird and mammal), geographical localization (each continent excepted Antarctica) and phylogenetic divergence/evolution (with the exclusion of rhabdovirus isolates presenting extremely large evolutionary distances). Sequences from this dataset, which encompassed virus isolates from 1955 to 2017, were download in November 2019 and were aligned using ClustalW 2.0 (Larkin et al., 2007). The most conserved regions were selected for primer design, based on the consensus sequence.

**Figure 2.**
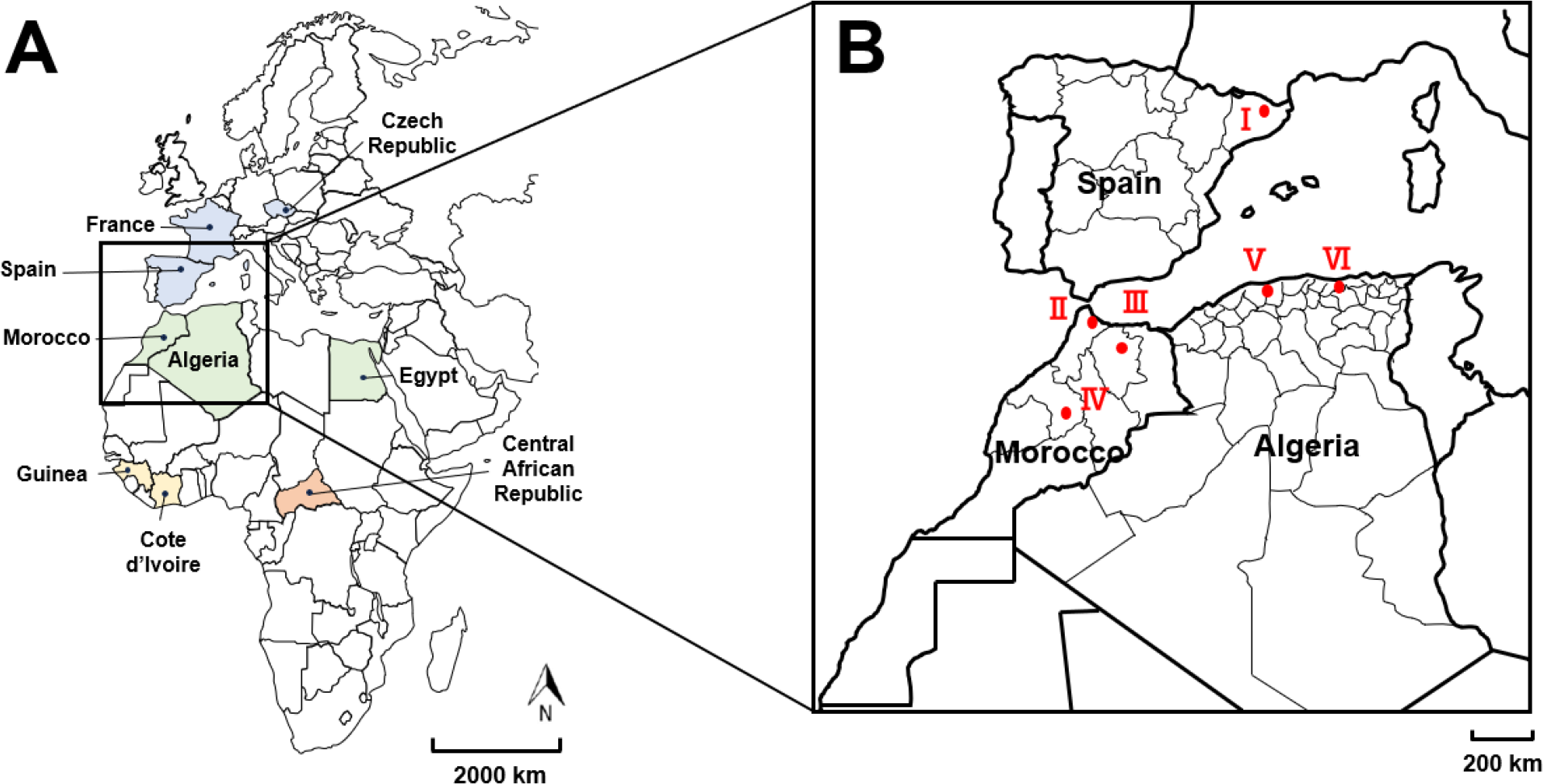
Geographical distribution of retrospective bat samples analyzed in this study. **A.** Geographical distribution at the country level, with samples collected from Europe (in blue) with Czechia, France and Spain; from North Africa (in green) with Algeria, Egypt and Morocco; from West Africa (in yellow) with Cote d’Ivoire and Guinea; and from central Africa (in orange) with Central African Republic. The distance scale is shown, as well as the North direction. **B.** Geographical distribution of the bat-related vesiculovirus MBV (Mediterranean bat virus) described in this study at the cave level, indicated with red dots and labelled by Roman numerals: cave I: A. Daví; cave II: Ghar- Knadel; cave III: Kef el Ghar; cave IV: Ifri N’Caid; cave V: Chrea; cave VI: Aokas. The distance scale is shown.

### 2.5 Pan-rhabdovirus PCR method validation

The determination of the optimal concentration of these primers was performed with serial dilutions (1:1, 1:10 and 1:100) of the reference strain Piry virus (PIRYV, BeAn 24232, 0413BRE) (*Vesiculovirus* genus) (Dacheux et al., 2010; De Souza, 2016). The primers were both tested by conventional PCR and SYBR Green-based real-time PCR (qPCR). Conventional PCR was performed with TaKaRa EX Taq kit (TaKaRa) and included 5 µL 10X PCR Buffer (Mg^2+^ plus), 4.0 µL dNTP Mix (2.5 mM each), 2.0 µL of both forward and reverse primers (20 µM or 200 µM), 2 µL of cDNA template and 5 µL of nuclease under 50 µL final volume. Real-time SYBR-based PCR (qPCR) was carried out with Power SYBR Green Master Mix kit (ThermoFisher) using 12.5 µL Mix Reagent, 1 µL of both forward and reverse primers (20 µM or 200 µM), 2 µL of cDNA template and completed to 25 µL with nuclease free water. Thermocycling conditions were performed according to the manufacturer’s instructions, and the annealing temperature (Tm) for each PCR system was calculated by TM calculator (http://www.Thermofisher.com).

The preliminary validation of the different pan-rhabdovirus screening assays was conducted on a limited panel (blood, brain and/or saliva) of negative and positive samples, including representative members of the genera *Ephemerovirus*, *Ledantevirus*, *Sunrhavirus* and *Vesiculovirus*. In addition, these methods were also compared to two previously published pan-rhabdovirus nested RT-PCR, with Pan- rhabdo RT-nPCR_1 (Aznar-Lopez et al., 2013) and Pan-rhabdo RT-nPCR_2 (Wray et al., 2016) (Table S3).

The sensitivity and specificity parameters of the selected pan-rhabdo molecular screening method (pan- rhabdo RT-nqPCR) was further validated using a large panel of positive and negative samples. The positive samples included representative virus members (n=60) of different genera (n=8) among the *Rhabdoviridae* family, and corresponded to laboratory or field samples (brain, blood or cell culture) from various animal species (*e.g.* bat, bovine, dog, fox, mouse) (Table 3). The negative samples encompassed viruses (*n* = 18) from other viral families (*n* = 9), as well as non-infected biological samples (*n* = 7) from different animal species (bat, dog, fox, monkey) and different types of tissues (brain, cell line) (Table 4). Based on this validation step, an analytic workflow based on the pan-rhabdo RT-nqPCR application was designed and applied for bat sample screening and interpretation.

### 2.6 Validation of bat positive samples obtained with the pan-rhabdo RT-nqPCR assay

Bat samples were considered positive for rhabdovirus when the characteristics of the dissociation curve (shape and melting temperature) observed with the amplicons were similar to those obtained with positive controls. Confirmation was done after Sanger sequencing of the amplicons and analysis of the sequences using Sequencher 5.2.4 (Gene Codes Corporation). In case of ambiguous results, amplicons were cloned using TOPO TA Cloning Kit (Invitrogen) according to the manufacturer’s instructions. In this case, a total of 5 to 10 clones for each sample was selected for Sanger sequencing, and sequence contigs were analyzed by BLASTn and BLASTx using the NCBI database.

### 2.7 Virus genome sequencing

Nearly complete genome sequences of bat-related rhabdoviruses were obtained using next generation sequencing (NGS) Illumina technology as previously described (Pallandre et al., 2020; Luo et al., 2021a). Contigs were obtained by combining *de novo* assembly and mapping (both with CLC Assembly Cell, Qiagen) using a dedicated workflow implemented on the Galaxy platform of Institut Pasteur (Mareuil, 2017; Pallandre et al., 2020; Luo et al., 2021a, 2021b). Contigs were then assembled and manually edited to produce the final consensus genome sequences using Sequencher. The quality and the accuracy of the final genome sequences were validated after a final mapping step of the original cleaned reads and visualized using Tablet (Milne et al., 2013).

Gaps or ambiguous nucleotide positions were corrected by (nested) PCR using TaKaRa EX Taq (TaKaRa) according to the manufacturer’s instructions, using specific primers designed from the genome sequences obtained by NGS (Table S4). Amplicons were submitted to Sanger sequencing and sequence correction was done using Sequencher.

Genome sequences were deposited in NCBI GenBank under the accession numbers MW491754- MW491760.

### 2.8 Genome analysis

Determination of the putative coding regions of the genome sequences was performed with Sequencher, and putative accessory genes was evaluated after comparison with similar rhabdovirus genome sequences available in GenBank.

Phylogenetic analysis of the bat vesiculoviruses and other members of the *Vesiculovirus* genus was conducted using a dataset of nucleotide sequences downloaded from NCBI GenBank (Table S5). For each virus, complete genome nucleotide sequences were aligned using ClustalW (version 2.0) (Larkin et al., 2007) and checked manually for accuracy. Phylogenetic trees were constructed with MEGA (version 7.0) (Kumar et al., 2016) and PhyML (version 3.0) (Guindon et al., 2010) using Maximum Likelihood method, with GTR+G+I model and with 1000 bootstrap replicates. Pairwise sequence identities were performed using MEGA.

### 2.9 Virus Isolation

Virus isolation was attempted using newborn suckling BALB/c mice (3 days old) (Charles River laboratories). For each positive sample, 20 to 50 µL lysed whole blood were diluted into 100 µL sterile PBS, gently mixed and centrifuged at 5000 g for 10 min at 4℃. Supernatant (5 to 10 µL) was inoculated intracerebrally to 4 to 5 newborn mice per sample. Inoculated animals were monitored daily and euthanized at day 5 or 7 after inoculation. Presence of virus was tested on brain after total RNA extraction and pan-rhabdo RT-nqPCR detection, as previously described.

## 3 Results

### 3.1 Primer design for the broad-spectrum detection of animal rhabdoviruses

A dataset of 103 nucleotide sequences of polymerases from rhabdovirus species belonging to 14 different genera were selected for multiple sequence alignment (Figure 2 and Table S2). After analysis, a highly conserved region between nucleotide positions 1600 and 2207 of the polymerase gene was selected for primer design (according to the sequence reference of Drosophila obscura Sigmavirus, DObSV, NCBI Accession NC_022580) (Figure 2). In total, 7 different primers were designed for 5 PCR systems, with 4 SYBR Green-based real-time PCR (qPCR) systems (screen-rhabdo qPCR_1 to screen-rhabdo qPCR_4) and 1 nested SYBR Green-based qPCR system (pan-rhabdo RT-nqPCR), the latter being also used as a nested conventional PCR (Table S3). The size of the different amplicons ranged from 119 to 575 pb.

### 3.2 Selection and validation of the primers for the broad-spectrum detection of animal rhabdoviruses

The pan-rhabdo RT-nqPCR system with a final primer concentration of 200 µM was selected using serial dilutions (1:1, 1:10 and 1:100) of the reference strain Piry virus (PIRYV, BeAn 24232, 0413BRE) (*Vesiculovirus* genus) and different primer concentrations (Table S6). A first screening test of the 7 different PCR assays was performed on a limited panel of positive samples (n=10) including 7 different rhabdovirus species divided into 4 field samples and 6 viruses isolated from inoculated mouse brains and 10 field negative controls (blood and saliva bat samples) (Table S7). The results obtained with the pan-rhabdo RT-nqPCR assay were all consistent, and this assay was selected for further validation.

### 3.3 Validation of the pan-rhabdo RT-nqPCR assay

Based on the preliminary validation step of the pan-rhabdo RT-nqPCR assay, a standardized process of analysis was established, based on two main parameters: Cp value and dissociation curve (Tm value and shape) (Figure S1). The sensitivity of this assay was evaluated using this process and a panel of 60 positive samples representative of 38 different rhabdovirus species among 8 different genera (Table 3). Among them, 21 were field samples from bat and dog brains or bovine blood. All these samples were detected with the pan-rhabdo RT-nqPCR assay, providing 100% sensitivity. The specificity of this assay was also evaluated on a panel of 18 viruses belonging to 9 different families other than *Rhabdoviridae*, and on 7 non-infected samples from 5 different animal species (Table 4). All these samples were not detected by the pan-rhabdo RT-nqPCR assay, exhibiting 100% specificity.

### 3.4 Bat sample screening with the pan-rhabdo RT-nqPCR assay

A total of 1962 bat samples, comprised of 423 saliva swabs, 816 blood and 723 brains, were tested with the pan-rhabdo RT-nqPCR assay. This large retrospective collection included samples from nine countries in Europe (Czechia, France and Spain) and Africa (Algeria, Central African Republic, Egypt, Guinea, Ivory Coast and Morocco) (Figure 1 and Table S1). These samples were collected from 2007 to 2019 and originated from 51 bat species, representative of 24 genera among eight different families, including frugivorous bat Pteropodidae and insectivorous bat Vespertilionidae, Miniopteridae, Rhinolophidae, Hipposideridae, Rhinopomatidae, Emballonuridae and Molossidae (Tables 1, 2 and S1).

**Table 1.** Distribution of the bat samples tested in this study according to the country of origin and to the sample type (saliva, blood pellet and brain).

| Region | Country | All Sample |  | Saliva sample |  | Blood sample |  | Brain sample |  |
| --- | --- | --- | --- | --- | --- | --- | --- | --- | --- |
|  |  | No. (%) | Species No. | No. (%) | Species No. | No. (%) | Species No. | No. (%) | Species No. |
| Europe | Czechia | 234 (12) | 5 | 0 | 0 | 0 | 0 | 234 (32.3) | 5 |
| Europe | France | 71 (3.6) | 9 | 0 | 0 | 0 | 0 | 71 (9.8) | 9 |
| Europe | Spain | 373 (19) | 5 | 341 (80.6) | 5 | 32 (4) | 2 | 0 | 0 |
| North Africa | Algeria | 232 (11.8) | 8 | 0 | 0 | 232 (28.4) | 8 | 0 | 0 |
| North Africa | Egypt | 148 (7.5) | 5 | 0 | 0 | 148 (18.1) | 5 | 0 | 0 |
| North Africa | Morocco | 420 (21.4) | 7 | 0 | 0 | 404 (49.5) | 7 | 16 (2.2) | 3 |
| West Africa | Ivory Coast | 96 (4.9) | 20 | 28 (6.6) | 13 | 0 | 0 | 68 (9.4) | 19 |
| West Africa | Guinee | 90 (4.6) | 8 | 54 (12.8) | 7 | 0 | 0 | 36 (5) | 7 |
| Central Africa | Central African Republic | 298 (15.2) | 6 | 0 | 0 | 0 | 0 | 298 (41.3) | 6 |
| **Total** |  | 1962 (100) |  | 423 (100) |  | 816 (100) |  | 723 (100) |  |

**Table 2.** Distribution of the bat samples tested in this study according to the bat families.

| Bat family | All sample |  | Saliva sample |  | Blood sample |  | Brain sample |  |
| --- | --- | --- | --- | --- | --- | --- | --- | --- |
|  | No. (%) | Species No. | No. (%) | Species No. | No. (%) | Species No. | No. (%) | Species No. |
| Pteropodidae | 514 (26.2) | 15 | 61 (14.4) | 7 | 100 (12.3) | 1 | 353 (48.8) | 14 |
| Emballonuridae | 18 (1) | 1 | 0 | 0 | 18 (2.2) | 1 | 0 | 0 |
| Hipposideridae | 40 (2) | 3 | 8 (1.9) | 3 | 0 | 0 | 32 (4.4) | 3 |
| Miniopteridae | 354 (18) | 1 | 193 (45.6) | 1 | 157 (19.2) | 1 | 4 (0.6) | 1 |
| Molossidae | 2 (0.1) | 1 | 0 | 0 | 0 | 0 | 2 (0.3) | 1 |
| Rhinolophidae | 183 (9.3) | 6 | 3 (0.7) | 2 | 164 (20.1) | 5 | 16 (2.2) | 3 |
| Rhinopomatidae | 28 (1.4) | 2 | 0 | 0 | 28 (3.4) | 2 | 0 | 0 |
| Vespertilionidae | 823 (42) | 22 | 158 (37.4) | 9 | 349 (42.8) | 7 | 316 (43.7) | 18 |
| Total | 1962 (100) | 51 | 423 (100) | 22 | 816 (100) | 17 | 723 (100) | 40 |

**Table 3.**
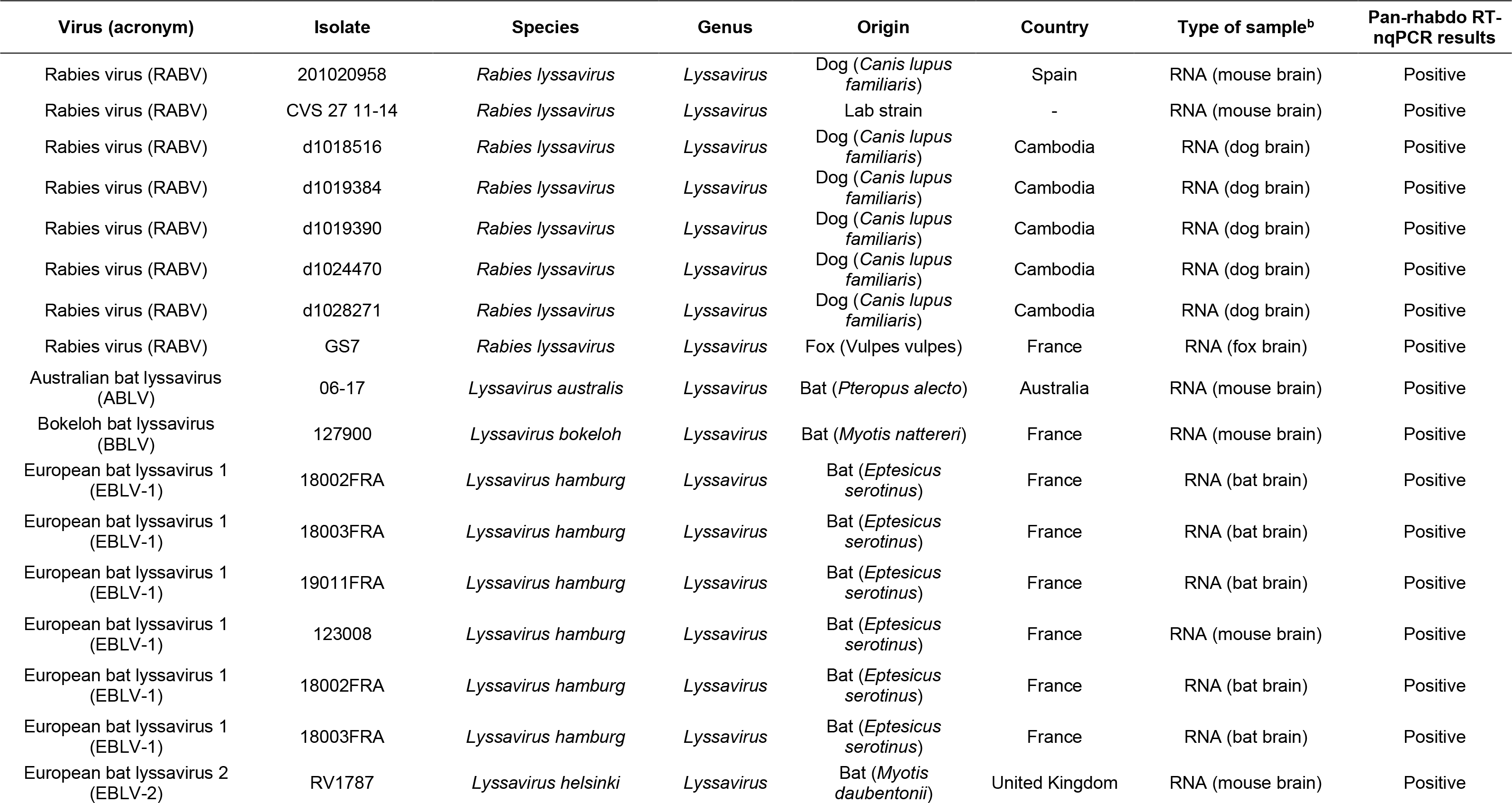
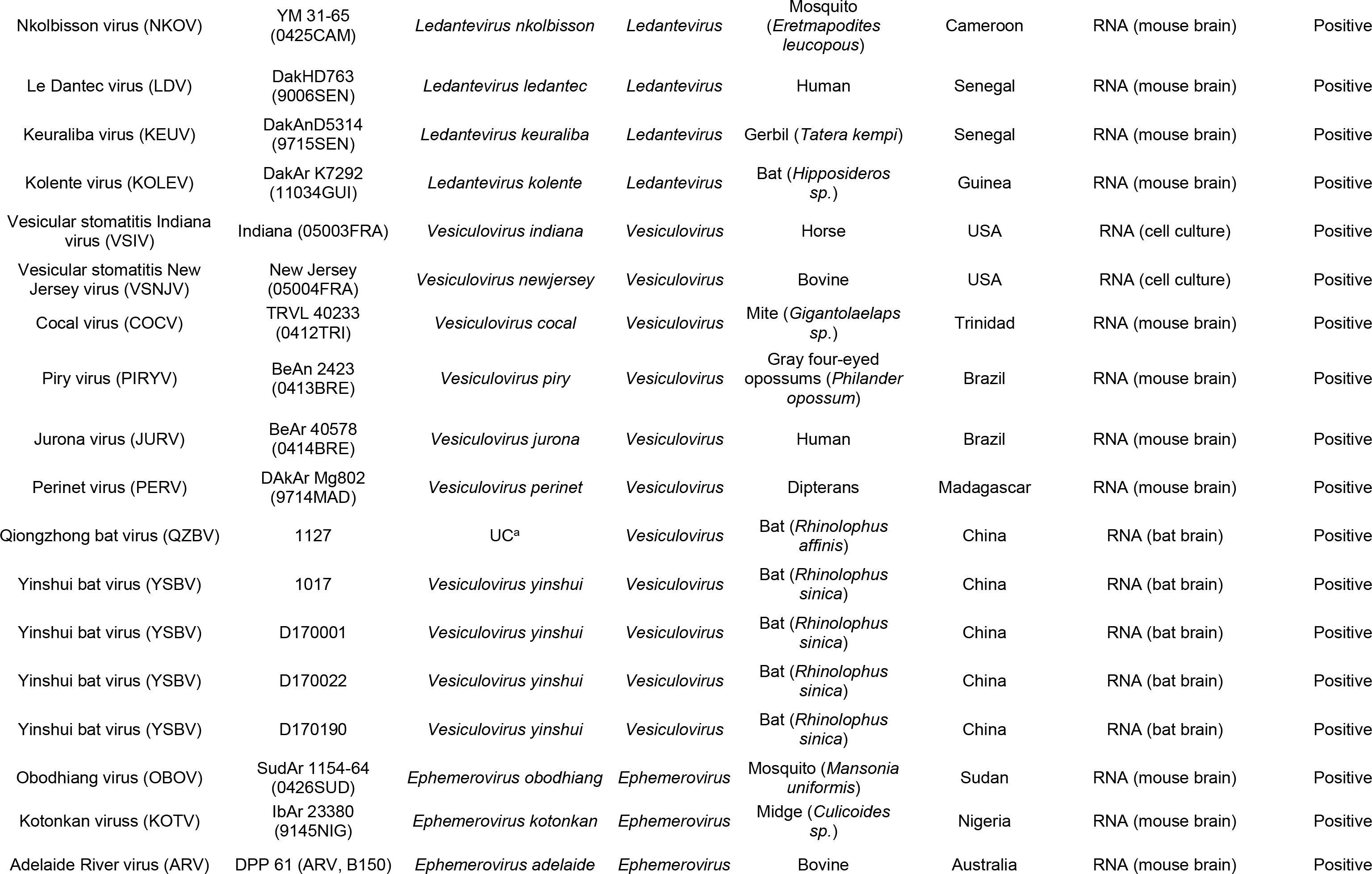
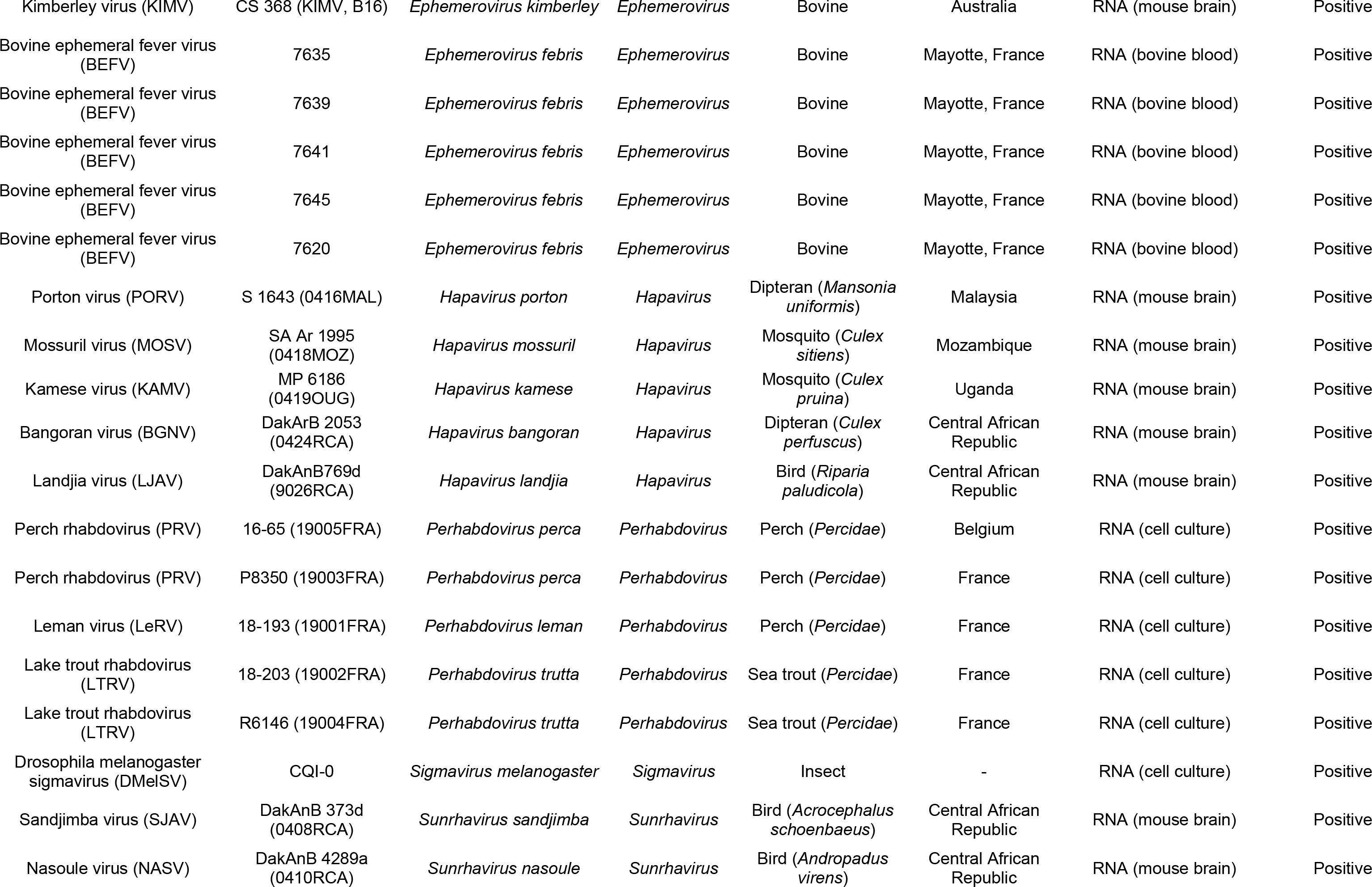
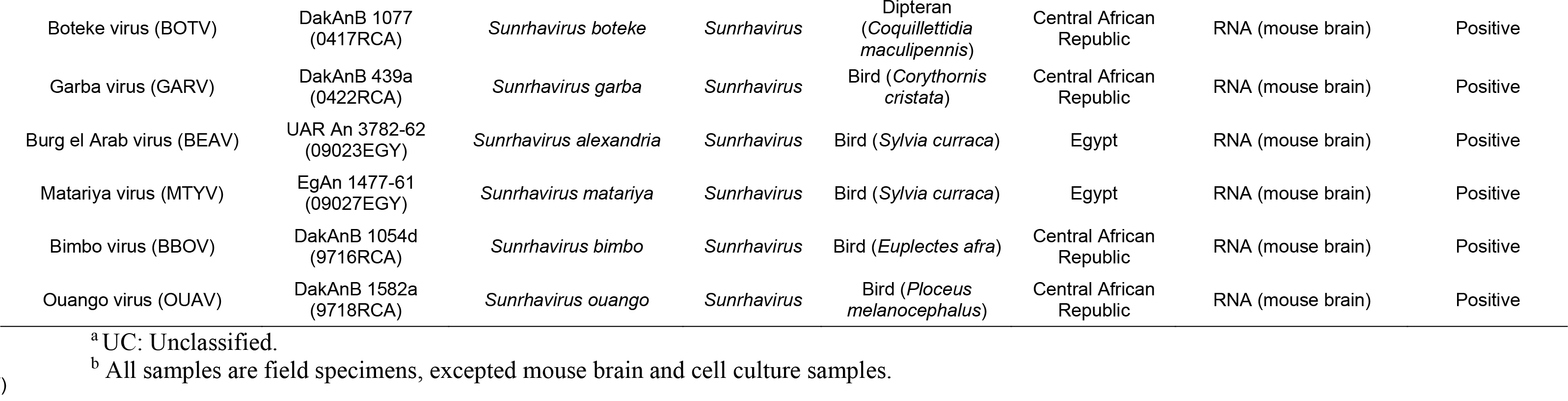
Detection results of the pan-rhabdo RT-nqPCR with a panel of representative members of different genera among the family *Rhabdoviridae*.

**Table 4.**
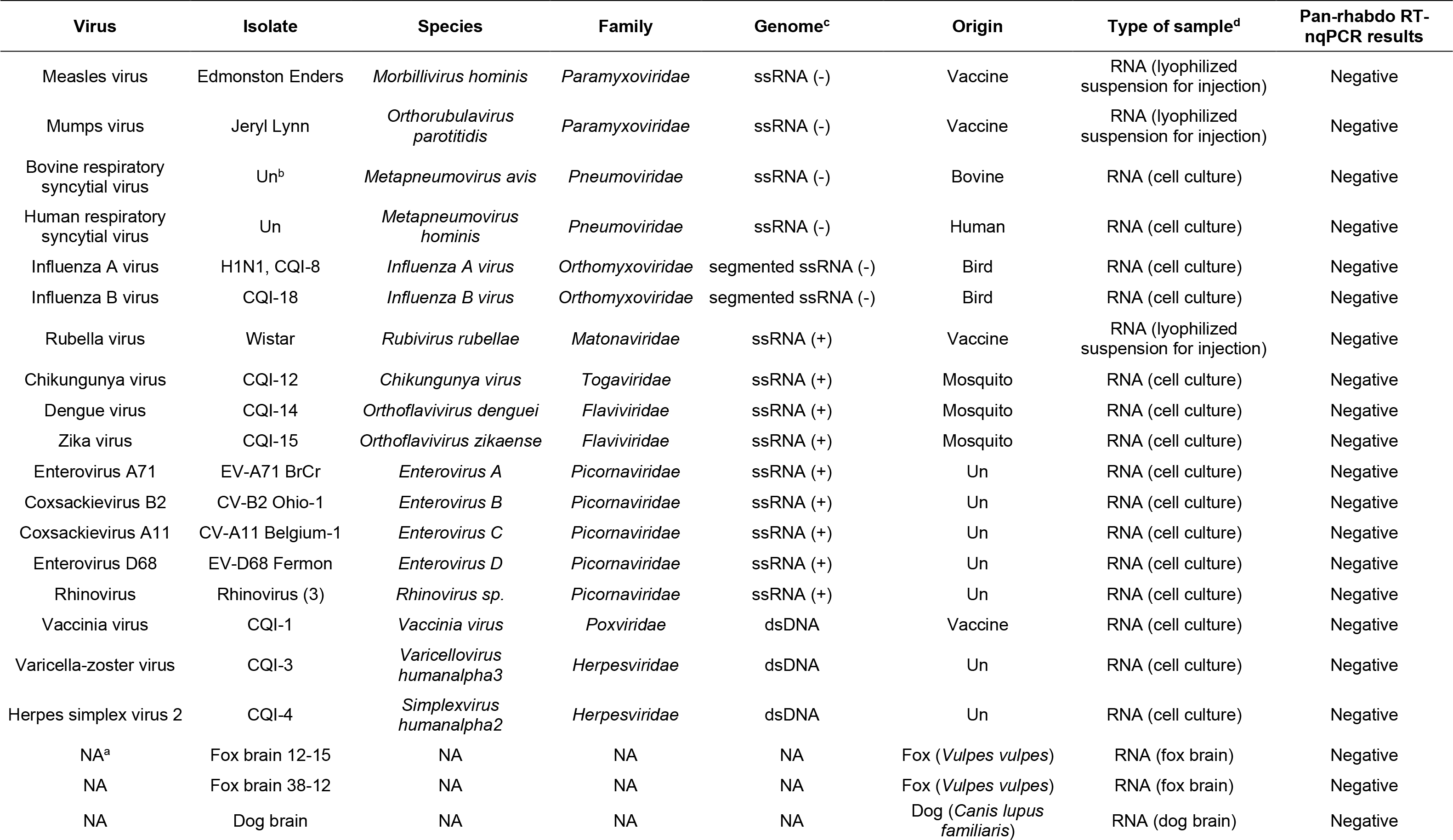
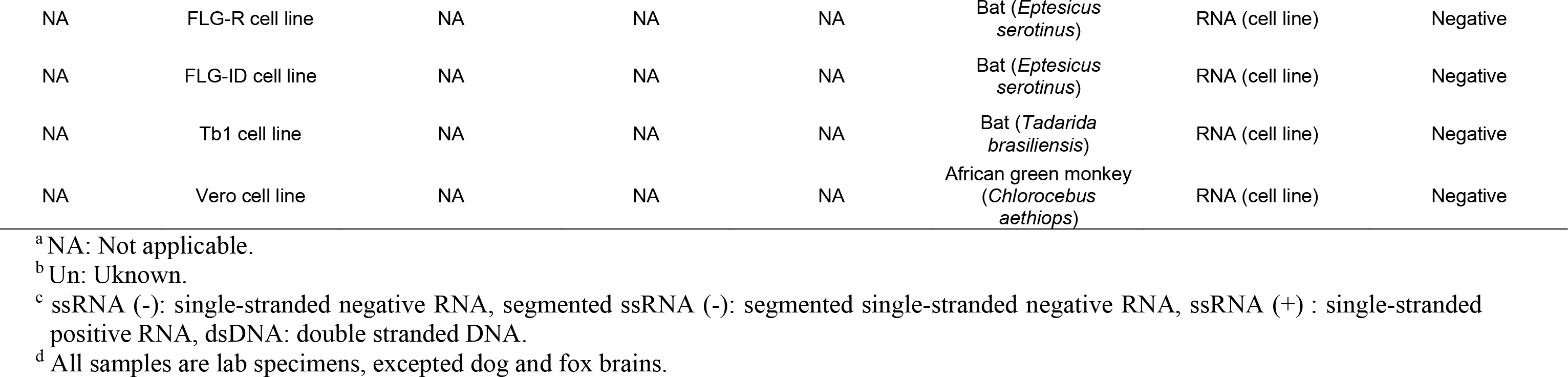
Specificity analysis of the pan-rhabdo RT-nqPCR on a panel of representative viruses other than rhabdoviruses and on a panel of non- infected samples.

| Virus | Isolate | Species | Family | Genome<sup>c</sup> | Origin | Type of sample<sup>d</sup> | Pan-rhabdo RT-nqPCR results |
| --- | --- | --- | --- | --- | --- | --- | --- |
| Measles virus | Edmonston Enders | *Morbillivirus hominis* | *Paramyxoviridae* | ssRNA (-) | Vaccine | RNA (lyophilized suspension for injection) | Negative |
| Mumps virus | Jeryl Lynn | *Orthorubulavirus parotitidis* | *Paramyxoviridae* | ssRNA (-) | Vaccine | RNA (lyophilized suspension for injection) | Negative |
| Bovine respiratory syncytial virus | Un<sup>b</sup> | *Metapneumovirus avis* | *Pneumoviridae* | ssRNA (-) | Bovine | RNA (cell culture) | Negative |
| Human respiratory syncytial virus | Un | *Metapneumovirus hominis* | *Pneumoviridae* | ssRNA (-) | Human | RNA (cell culture) | Negative |
| Influenza A virus | H1N1, CQI-8 | *Influenza A virus* | *Orthomyxoviridae* | segmented ssRNA (-) | Bird | RNA (cell culture) | Negative |
| Influenza B virus | CQI-18 | *Influenza B virus* | *Orthomyxoviridae* | segmented ssRNA (-) | Bird | RNA (cell culture) | Negative |
| Rubella virus | Wistar | *Rubivirus rubellae* | *Matonaviridae* | ssRNA (+) | Vaccine | RNA (lyophilized suspension for injection) | Negative |
| Chikungunya virus | CQI-12 | *Chikungunya virus* | *Togaviridae* | ssRNA (+) | Mosquito | RNA (cell culture) | Negative |
| Dengue virus | CQI-14 | *Orthoflavivirus denguei* | *Flaviviridae* | ssRNA (+) | Mosquito | RNA (cell culture) | Negative |
| Zika virus | CQI-15 | *Orthoflavivirus zikaense* | *Flaviviridae* | ssRNA (+) | Mosquito | RNA (cell culture) | Negative |
| Enterovirus A71 | EV-A71 BrCr | *Enterovirus A* | *Picornaviridae* | ssRNA (+) | Un | RNA (cell culture) | Negative |
| Coxsackievirus B2 | CV-B2 Ohio-1 | *Enterovirus B* | *Picornaviridae* | ssRNA (+) | Un | RNA (cell culture) | Negative |
| Coxsackievirus A11 | CV-A11 Belgium-1 | *Enterovirus C* | *Picornaviridae* | ssRNA (+) | Un | RNA (cell culture) | Negative |
| Enterovirus D68 | EV-D68 Fermon | *Enterovirus D* | *Picornaviridae* | ssRNA (+) | Un | RNA (cell culture) | Negative |
| Rhinovirus | Rhinovirus (3) | *Rhinovirus sp.* | *Picornaviridae* | ssRNA (+) | Un | RNA (cell culture) | Negative |
| Vaccinia virus | CQI-1 | *Vaccinia virus* | *Poxviridae* | dsDNA | Vaccine | RNA (cell culture) | Negative |
| Varicella-zoster virus | CQI-3 | *Varicellovirus humanalpha3* | *Herpesviridae* | dsDNA | Un | RNA (cell culture) | Negative |
| Herpes simplex virus 2 | CQI-4 | *Simplexvirus humanalpha2* | *Herpesviridae* | dsDNA | Un | RNA (cell culture) | Negative |
| NA<sup>a</sup> | Fox brain 12-15 | NA | NA | NA | Fox (*Vulpes vulpes*) | RNA (fox brain) | Negative |
| NA | Fox brain 38-12 | NA | NA | NA | Fox (*Vulpes vulpes*) | RNA (fox brain) | Negative |
| NA | Dog brain | NA | NA | NA | Dog (*Canis lupus familiaris*) | RNA (dog brain) | Negative |
| NA | FLG-R cell line | NA | NA | NA | Bat (*Eptesicus serotinus*) | RNA (cell line) | Negative |
| NA | FLG-ID cell line | NA | NA | NA | Bat (*Eptesicus serotinus*) | RNA (cell line) | Negative |
| NA | Tb1 cell line | NA | NA | NA | Bat (*Tadarida brasiliensis*) | RNA (cell line) | Negative |
| NA | Vero cell line | NA | NA | NA | African green monkey (*Chlorocebus aethiops*) | RNA (cell line) | Negative |

<sup>a</sup> NA: Not applicable.
<sup>b</sup> Un: Unknown.
<sup>c</sup> ssRNA (-): single-stranded negative RNA, segmented ssRNA (-): segmented single-stranded negative RNA, ssRNA (+) : single-stranded positive RNA, dsDNA: double stranded DNA.
<sup>d</sup> All samples are lab specimens, excepted dog and fox brains.

Among the 1962 samples, 23 (1.2%) were found positive for rhabdovirus detection. All of them originated from 3 different bat species, including 6 samples from *Miniopterus schreibersii*, 2 from *Rhinolophus euryale* and 15 from *Rhinolophus ferrumequinum*, leading to a prevalence of 1.7% (*n* = 354), 4.9% (*n* = 41) and 11.9% (*n* = 126), respectively, when considering the total number of samples tested for each of these species (Table 5). All the positive bats originated from the Mediterranean region, with 6 bats from Spain, 11 from Algeria and 6 from Morocco (Table 5), resulting in a prevalence per country of 1.6% (*n* = 373), 4.7% (*n* = 232) and 1.4% (*n* = 420), respectively. None of the brain samples tested positive, whereas the detection of rhabdovirus by the pan-rhabdo RT-nqPCR assay was obtained for 6 saliva samples (1.4%, *n* = 423) and 17 blood samples (2%, *n* = 816), collected over the years 2008, 2009 and 2012 (Table 6). Sanger sequencing of the amplicons confirmed the detection of rhabdoviruses, probably related to *Vesiculovirus* genus.

**Table 5.**
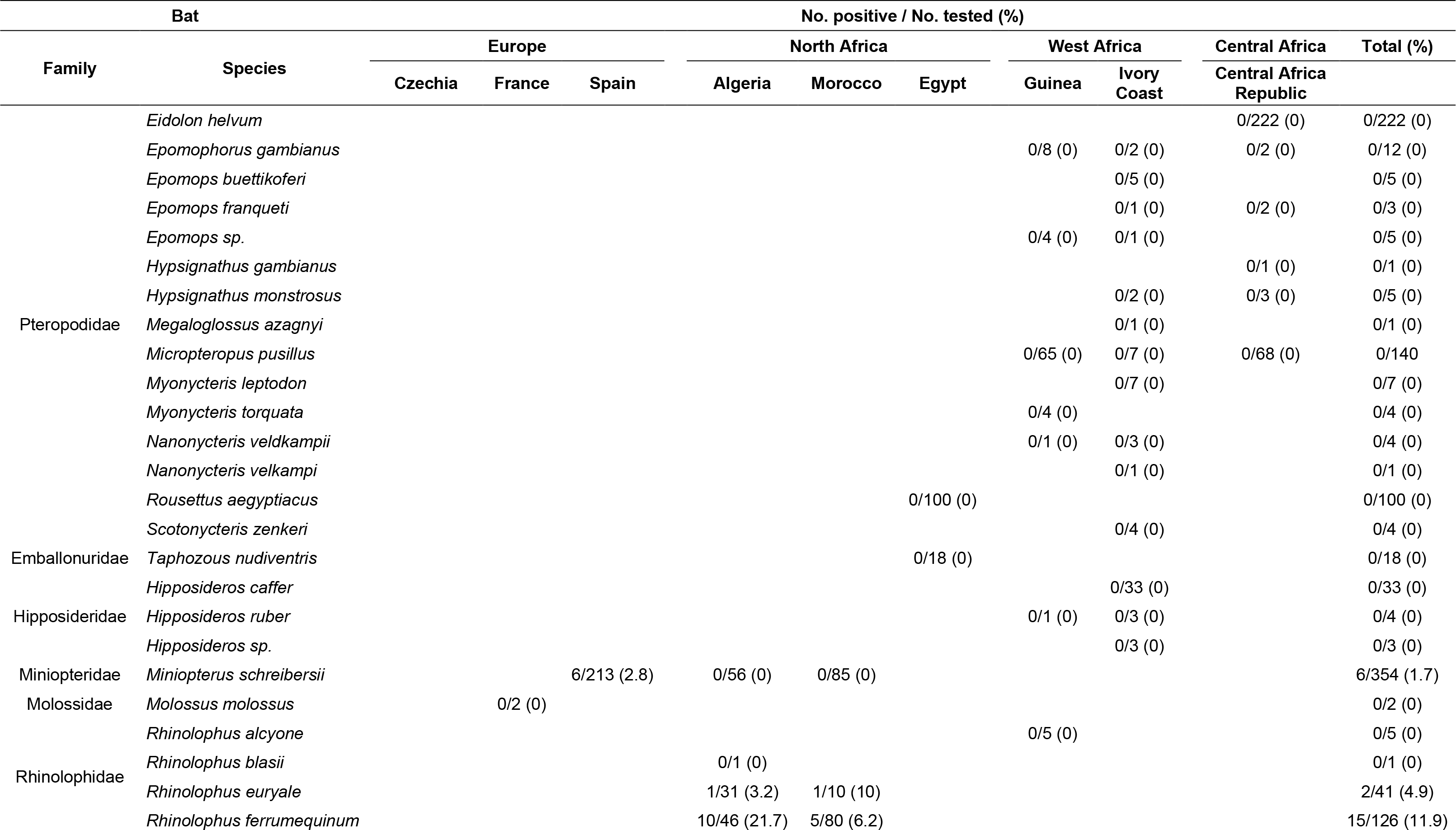
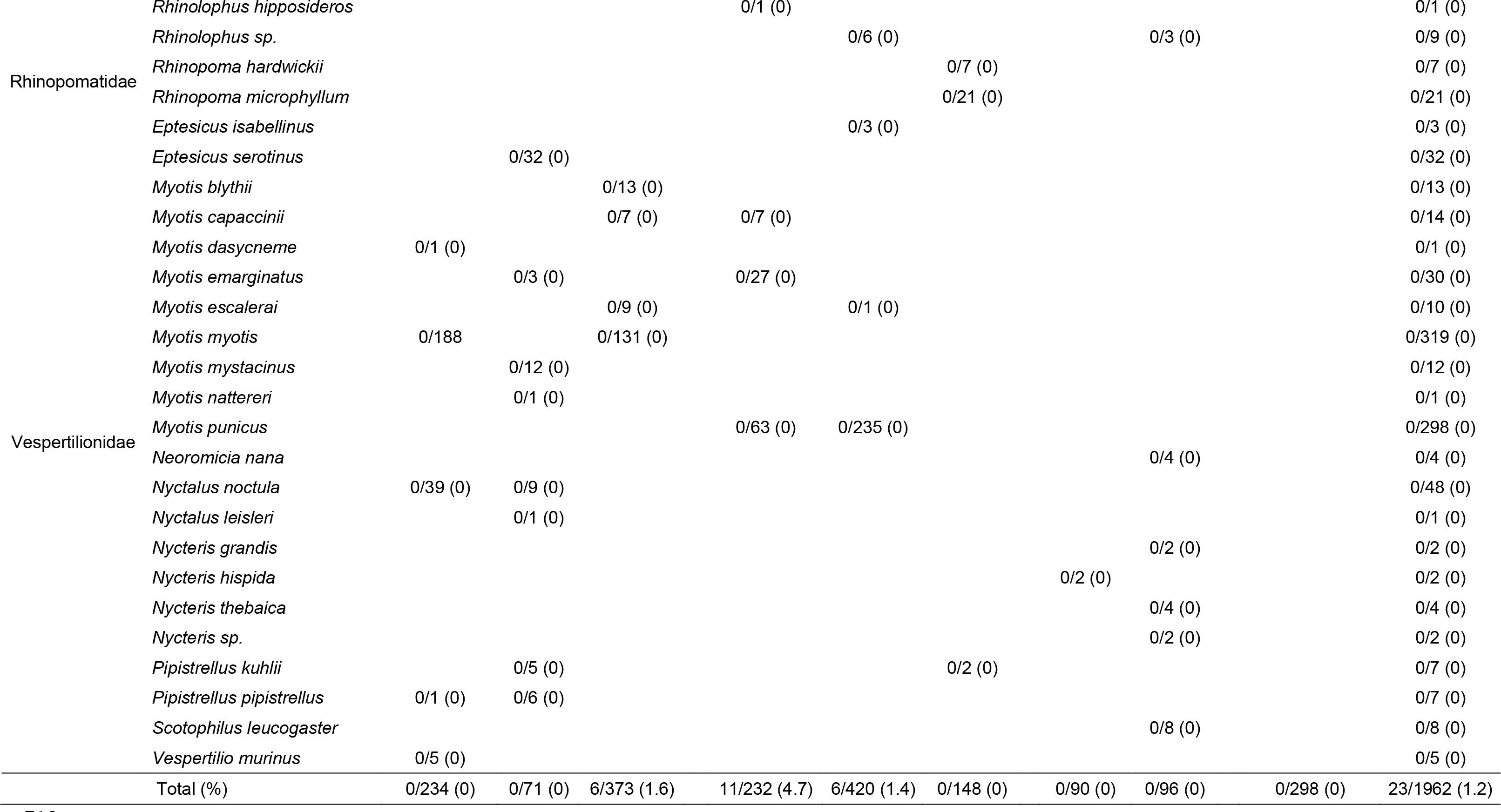
Results of the rhabdovirus detection with the pan-rhabdo RT-nqPCR for a collection of retrospective bat samples (blood, brain, saliva) according to bat species and geographical location.

**Table 6.** Results of the detection rate of rhabdovirus with the pan-rhabdo RT-nqPCR in saliva and brain samples collected from different bat species and in selected caves from Algeria, Morocco and Spain.

| Colletion<br>year | No. positive / No. tested (%) |  |  |  |  |  |  | Total (%) |
| --- | --- | --- | --- | --- | --- | --- | --- | --- |
|  | *Miniopterus schreibersii* | *Rhinolophus euryale* |  | *Rhinolophus ferrumequinum* |  |  |  |  |
|  | Cave<sup>a</sup> I (Spain) | Cave III<br>(Morocco) | Cave VI<br>(Algeria) | Cave II<br>(Morocco) | Cave IV<br>(Morocco) | Cave V<br>(Algeria) | Cave VI<br>(Algeria) |  |
| 2008 |  | 1/4 (25) | 1/28 (3.6) | 3/40 (7.5) |  | 1/7 (14.3) |  | 6/79 (7.6) |
| 2009 |  |  |  |  | 2/14 (14.3) | 7/29 (24.1) | 2/5 (40) | 11/48 (22.9) |
| 2012 | 6/18 (33.3) |  |  |  |  |  |  | 6/18 (33.3) |
| Total (%) | 6/18 (33.3) | 1/4 (25) | 1/28 (3.6) | 3/40 (7.5) | 2/14 (14.3) | 8/36 (22.2) | 2/5 (40) | 23/145 (15.9) |

<sup>a</sup> The identification of the caves is as follows: cave I: A. Daví, cave II: Ghar-Knadel, cave III: Kef el Ghar, cave IV: Ifri N'Caid, cave V: Chrea, cave VI: Aokas.

All the *Miniopterus schreibersii* positive samples were found in saliva in Spain in the Catalonian cave I (A. Daví) from 2012, which corresponded to a prevalence of 33.3% of the total saliva number tested from this cave (n=18) (Figure 2) (Table 6). The two positive samples from *Rhinolophus euryale* were detected in blood pellet from cave III and cave VI in Morocco and Algeria from 2008, respectively. The positive rate in blood for this bat species was 25% (*n* = 4) and 3.6 (*n* = 28) for cave III and VI, respectively (Table 6). Lastly, the highest number of positive samples (*n* = 15) was found in the blood pellet of *Rhinolophus ferrumequinum* bats collected in two caves (caves II and IV) in Morocco and in two caves (caves V and VI) in Algeria during the period of 2008-2009 (Figure 2) (Table 6). The overall prevalence in each of these caves for this bat species was 7.5% (*n* = 40), 14.3% (*n* = 14), 22.2% (*n* = 36) and 40% (*n* = 5) for caves II, IV, V and VI, respectively. Interestingly, samples from cave V in Algeria were collected both in 2008 and 2009, with a prevalence of 14.3% (*n* = 7) and 24.1% (*n* = 29), respectively. All the bat species and other samples available and tested from these six caves were negative.

### 3.5 Complete genome analysis of bat rhabdoviruses

Between 3 to 14 million raw reads were obtained per sample (around 5 million reads on average) for all the 23 positive samples. Specific reads or contigs for rhabdovirus were obtained for all of them, but only 16 samples exhibited a large genome contigs after analysis (Table S8). The average coverage for each of them varied from 1.58x to 2472x. The remaining gaps and low coverage regions were resolved through specific PCR or nested-PCR, following by Sanger sequencing of the corresponding amplicons. After a last mapping round using the sequence consensus, 16 nearly complete genome sequences were obtained, ranged from 10,914 to 11,093 nt in length (excluding 3′ Leader and 5′ Trailer sequences (Table 7).

**Table 7.** Reads coverage of the 16 Mediterranean bat virus (MBV) isolates genome sequences after the last mapping step using NGS data.

| Isolate | Country | Sample type | Raw reads<br>(No. x10<sup>6</sup>) | Reads cleaned<br>(No. x10<sup>6</sup>) | Mapped<br>reads (No.) | Sequence<br>size (nt) | Average<br>coverage (X) | Length sequence<br>coverage (%) |
| --- | --- | --- | --- | --- | --- | --- | --- | --- |
| A08011 | Algeria | Blood | 3.798 | 2.662 | 1877 | 10,914 | 25.35 | 100 |
| A08065 | Algeria | Blood | 3.842 | 2.735 | 1156 | 10,938 | 15.71 | 100 |
| A09061 | Algeria | Blood | 5.152 | 4.057 | 189,052 | 11,092 | 2472.19 | 100 |
| A09097 | Algeria | Blood | 6.270 | 4.734 | 11,433 | 11,092 | 151.80 | 100 |
| A09145 | Algeria | Blood | 3.580 | 2.596 | 57,269 | 11,093 | 758.85 | 100 |
| A09151 | Algeria | Blood | 5.555 | 4.141 | 15,805 | 11,093 | 210.67 | 100 |
| A09153 | Algeria | Blood | 3.415 | 2.447 | 19,037 | 11,065 | 251.75 | 100 |
| A09181 | Algeria | Blood | 4.662 | 3.452 | 128,719 | 11,093 | 1686.87 | 100 |
| A09193 | Algeria | Blood | 3.398 | 2.543 | 4658 | 11,093 | 61.87 | 100 |
| A09197 | Algeria | Blood | 6.372 | 4.938 | 2464 | 11,011 | 33.09 | 100 |
| M08013 | Morocco | Blood | 5.570 | 3.889 | 3233 | 10,980 | 43.46 | 100 |
| M08017 | Morocco | Blood | 3.854 | 2.743 | 3740 | 10,932 | 50.79 | 100 |
| M08051 | Morocco | Blood | 5.391 | 3.860 | 1085 | 10,975 | 14.65 | 100 |
| M09005 | Morocco | Blood | 4.787 | 3.793 | 4278 | 10,920 | 58.21 | 100 |
| M09009 | Morocco | Blood | 5.006 | 3.893 | 38,266 | 11,039 | 513.67 | 100 |
| 2012096 | Spain | Saliva | 14.188 | 9.189 | 122 | 11,068 | 1.58 | 52.2 |

The 16 genome sequences exhibited a common and typical rhabdovirus organization, consisting of five canonical genes encoding, in the following order, the nucleoprotein (N) (422 aa, 1269 nt), the phosphoprotein (P) (226 aa, 681 nt), the matrix protein (M) (205 aa, 618 nt), the glycoprotein (G) (514 aa, 1545 nt), and the RNA polymerase (L) (2102 aa, 6309 nt) (Figure 3A). For 4 genome sequences (A08011, A08065, M08017 and M09005), the N coding sequence remained incomplete after sequencing, with 40-60 missing nucleotides after the start codon. The transcription initiation (TI) signal was highly conserved, with the TAGCAGRG sequence (R=A/G), whereas the transcription termination (TTP) signal was YATG(A)7 consensus sequence (Y=C/T) (Figure 3B). In addition, two potential coding sequences representing putative accessory genes U1 (73 aa, 222 nt) and U2 (63 aa, 192 nt) were identified within the G gene (Figure 3A). After concatenation of the coding sequences for each genome and pairwise comparison, the sequences exhibited high nucleotide identities, with 97.3 % to 99.9 %, suggesting that all of them were classified into one species tentatively named Mediterranean bat virus (MBV) (Table S9).

**Figure 3.**
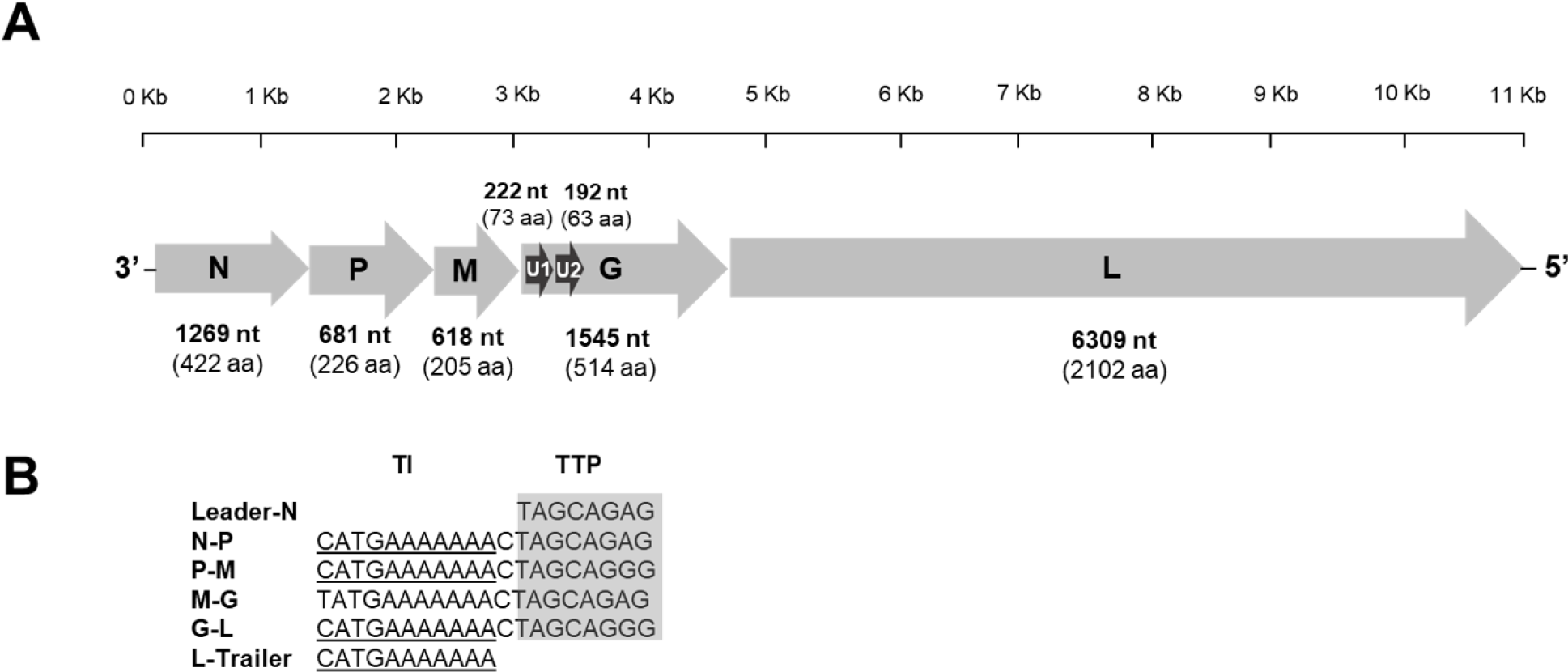
Analysis of the genome of the Mediterranean bat virus (MBV) detected by pan-rhabdo RT- nqPCR. **A.** Schematic organization of the genome. Grey arrows represent the five canonical open reading frames (ORFs) with the nucleoprotein (N), the phosphoprotein (P), the matrix protein (M), the glycoprotein (G) and the RNA polymerase (L). Black arrows indicate the position of putative additional ORFs. The nucleotide and amino acid lengths of each ORF are indicated. **B.** Description of the transcription initiation (TI) and the transcription termination (TTP) signal sequences. Conserved motifs are indicated in grey for TI or underlined for TTP. The consensus sequences are indicated, with Y = C/T, M = A/C, W = C/G and R = A/G.

Lastly, BLASTn and BLASTx analyses performed on the 16 genome sequences demonstrated that all these viruses were most closely related to bat rhabdoviruses belonging to the genus *Vesiculovirus*, and more particularly to Chinese bat-related rhabdoviruses recently described, such as Yinshui bat virus (GenBank number MN607597) or Qiongzhong bat virus (GenBank number MN607593) (Luo et al., 2021a).

Similar sequence comparison and BLAST analyses conducted on the short contigs (< 300 nt) obtained after NGS with the 7 other positive bat samples demonstrated that each of them was closely related to MBV.

### 3.6 Phylogenetic analysis of the Mediterranean bat virus

A maximum-likelihood phylogenetic analysis was performed on the 16 genome nucleotide sequences of MBV and representative members of vesiculoviruses. The phylogeny confirmed that MBV belonged to the *Vesiculovirus* genus. Among this genus, three distinct phylogroups were identified, with the phylogroup I clustered various viruses from America whereas the phylogroup II was cosmopolitan, encompassing viruses from Africa, American, Asia and Europe (Figure 4). Interestingly, all bat-related vesiculoviruses were clustered into the same phylogroup III. Within this phylogroup, a geographical clustering was observed, with three main groups: one encompassing the bat vesiculoviruses obtained from *Eptesicus fuscus* in North America (with ABV), the second one grouping bat vesiculovirus from Asia (and more specifically China) with two distinct branches according to the host bat species (*Rhinolophus affinis* or *Rhinolophus. sinicus*), and the last one clustering together all the MBV isolates from the Mediterranean region, regardless of the host bat species (Figure 4).

**Figure 4.**
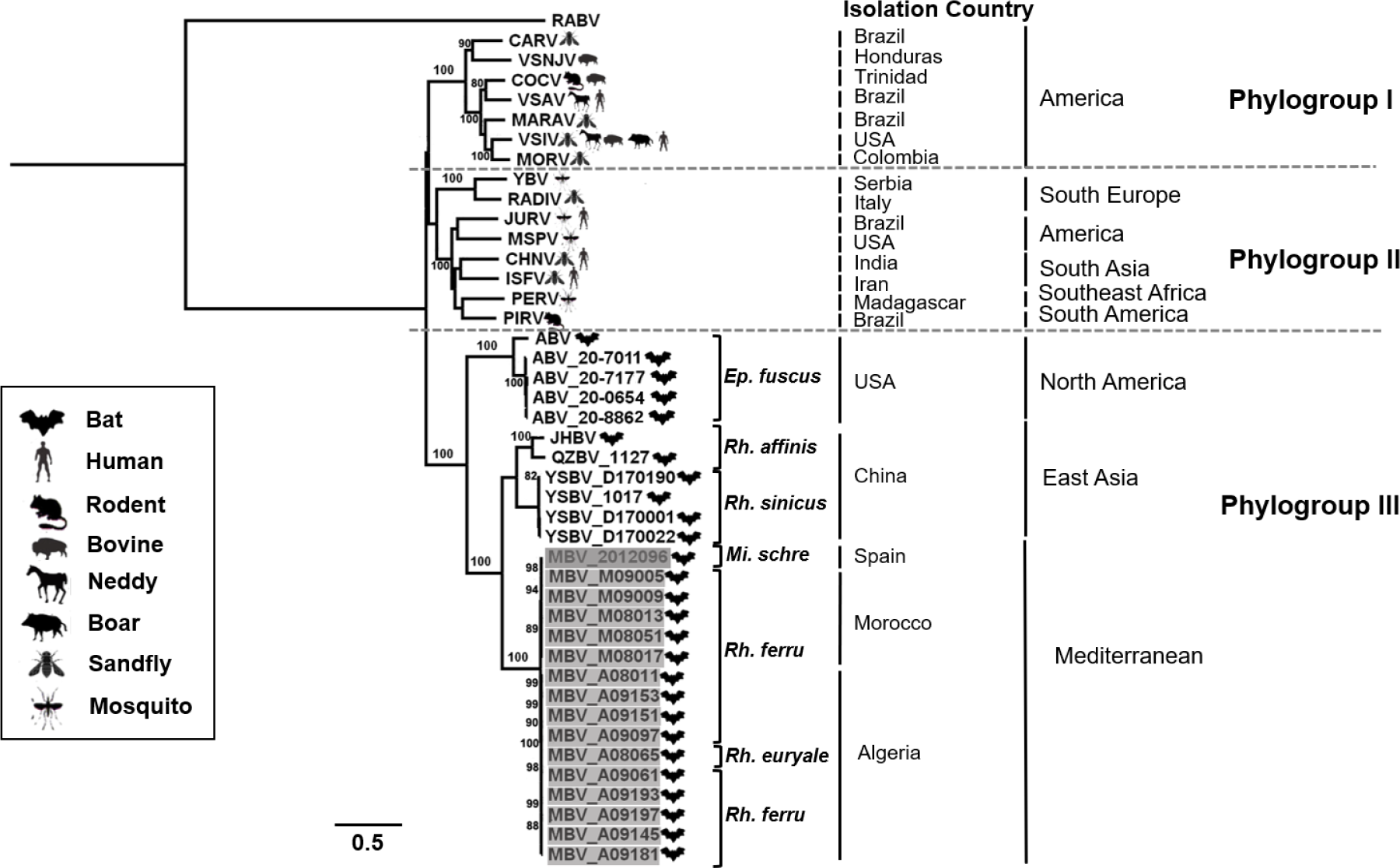
Phylogenetic classification of all members of the genus *Vesiculovirus,* including the Mediterranean bat virus (MBV) detected in this study. A maximum likelihood phylogenetic tree was done with PhyML3.0 on the nucleotide of complete genome, using the GTR+G+I model and with 1000 bootstrap replicates. The main animal reservoirs for each virus are indicated by specific cartoons and the 16 bat rhabdovirus isolates of MBV described in this study are indicated in grey. The bat species information was represented by abbreviation on the figure (*Ep.fuscus: Eptesicus fuscus, Rh.affinis: Rhinolophus affinis, Rh.sinicus: Rhinolophus sinicus, Mi.schre: Miniopterus schreibersii, Rh.ferru: Rhinolophus ferrumequinum, Rh.euryale: Rhinolophus euryale*), and the isolation country for each virus was presented in the right of the illustration. All bootstrap proportion values (BSP) > 80% are specified. Scale bar indicates nucleotide substitutions per site. The classical rabies virus (RABV) was included for out clade in the phylogenetic analysis.

### 3.7 Genetic diversity of the Mediterranean bat virus

A more detailed genetic analysis was performed for the available 16 genome sequences of MBV, in order to compare the different isolates according to their geographical origin and/or host bat species. At the scale of the phylogenetic analysis carried out on the genomic sequences, a clustering is observed between the country of origin, and more precisely between the location of the caves. For example, Algerian MBV isolates formed a distinct and well supported group, subdivided into subclusters according to the cave origins, i.e. cave VI, V and VI (Figure 5). Two phylogroups were also identified for the Moroccan strains, depending on the cave origin (cave II and cave IV). Interestingly, the single Spanish MBV isolate was found to be more closely related to the MBV viruses identified in the cave IV from Morocco, while being genetically distinct from it (Figure 5).

**Figure 5.**
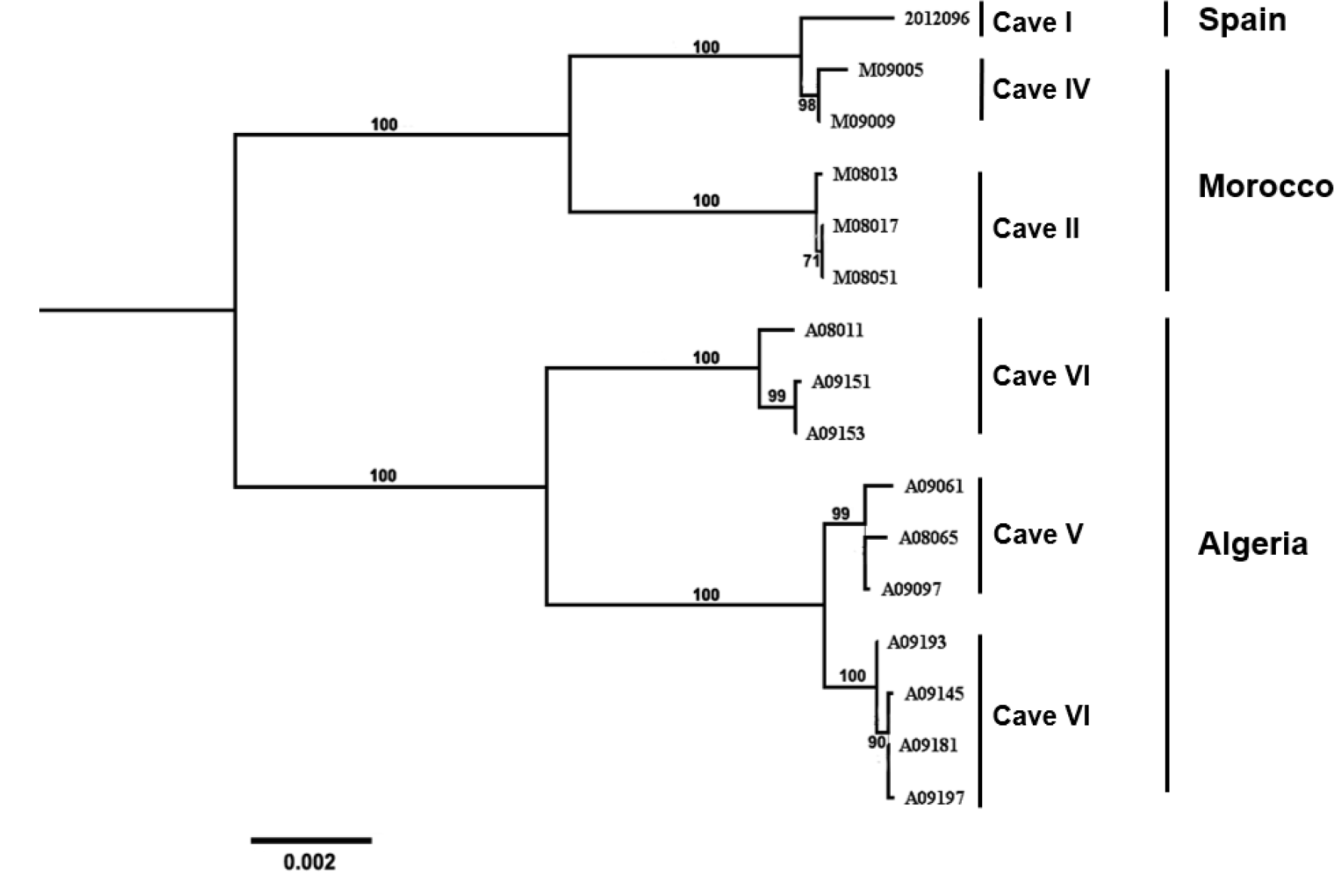
Details of the phylogenetic relationship between the 16 isolates of Mediterranean bat virus (MBV). A maximum likelihood phylogenetic tree was done with PhyML3.0 on the nucleotide of complete genome, using the GTR+G+I model and with 1000 bootstrap replicates. All bootstrap proportion values (BSP) > 70% are specified. Scale bar indicates nucleotide substitutions per site. The identification of the caves and of the country of origin is indicated.

At the overall genome sequence, the nucleotide identity was very high between the 16 sequences of MBV isolates, ranging from 97.3 to 99.9% (Table S9). The deducted amino acid identity of each five canonical proteins was also high between the 16 genome sequences of MBV isolates, with 100% identities for the N and M proteins, between 98.6 % and 100 % for the P and G proteins, and ≥ 99% for the L proteins (Table S10). These amino acid identity results indicated that the 16 different MBV isolates were related to the same virus species.

Within the genus *Vesiculovirus*, the highest percentage of deducted amino-acid identity for each of the 5 coding regions (N, P, M, G and L) was observed for the species *Vesiculovirus yinshui* (with isolate YSBV 1017), following with the species *Vesiculovirus jinghong* (with isolates QZBV 1127 and JHBV) (Table S11). For the first and more closely related species YSBV, these values were 75.5%, 21.1%, 61.5%, 62.4% and 67% for the N, P, M, G and L proteins, respectively (Table S11). These results suggest that MBV virus represents a novel vesiculovirus species.

Among the P, G and L deducted proteins, a total of 28 mutations were found: 2 for P, 7 for G and 19 for L proteins. As expected, the number of amino acids mutations was lower within MBV belonging to the same geographical cluster. A total of 8 mutations was found for the 10 isolates from Algeria, with one in the G proteins and 7 in the L proteins, similarly, whereas 7 mutations were observed within the 5 viruses from Morocco, with 2 in the P protein, 2 in the G protein 3 in the L protein (Figure S2). As expected from the phylogenetic analysis, the Spanish strain was closer to the Moroccan strains, with only 1 mutation in the G protein and 4 in the L protein with the strain M09009 from cave IV.

### 3.8 Virus Isolation

Virus isolation was performed after intracranial inoculation of newborn BALB/c mice (3 days old) with remaining blood pellets. After 7 days of observation, no clinical symptoms neither mortality was observed. Results obtained with the pan-rhabdo-RT-qPCR after total RNA of the inoculated mice brains remained negative.

## 4 Discussion

Among the various viral families which form the virosphere, the *Rhabdoviridae* family is one of the largest, comprising nearly 320 different species in 46 defined genera and 3 subfamilies (Kuhn et al., 2022; Walker et al., 2022b) (https://talk.ictvonline.org/). So far, at least 30 distinct rhabdovirus species belonging to four genera - *Lyssavirus*, *Vesiculovirus*, *Ledantevirus* and *Alphanemrhavirus* - were found in bats (Kuhn et al., 2022; Walker et al., 2022b). Indeed, nearly a hundred species of insectivorous and frugivorous bats were identified as infected or exposed to rhabdoviruses, mainly lyssaviruses (http://www.mgc.ac.cn/DBatVir/). However, other bat-related rhabdoviruses were also detected, but to a lesser extent. Given the extreme diversity of bat species (> 1400 different species) and their ability to host a high number of viruses, it is very likely that the diversity of rhabdoviruses hosted in these animals remains largely unknown.

In order to explore more in details this diversity and the rhabdovirus prevalence in bats, we successfully developed and validated a combined nested RT-qPCR technique (pan-rhabdo RT-nqPCR) dedicated to the broad detection of animal rhabdoviruses and targeting a conserved region in the polymerase gene. The comparison of this technique with similar previously published generic methods, such as the nested DimLis PCR RT-PCR (named “Pan-rhabdo RT-nPCR_1” in our study) (Aznar-Lopez et al., 2013) or the RT-PCR based on primers PVO3/PVO4/ PVOnstF (identified as “Pan-rhabdo RT-nPCR_2” in our study) (Wray et al., 2016), suggested a higher sensitivity, although the sample panel used was limited.

The pan-rhabdo RT-nqPCR was used for the screening of a large collection of retrospective bat samples originated from different countries. The overall prevalence was low, with 23 samples (1.2%) positive in total, including 17 blood pellet (2.1%) and 6 saliva (1.4%) samples. Only three out the 51 different bat species tested were found positive, with *Miniopterus schreibersii* (n=6), *Rhinolophus euryale* (n=2) and *Rhinolophus ferrumequinum* (n=15). Interestingly, all the positive bats were collected during 2008- 2012 in the Mediterranean basin, with Algeria (n=11) and Morocco (n=6) for the two *Rhinolophus* species, and Spain (n=6) for the *Miniopterus* species.

Although low, this prevalence rate is higher that found in other studies. For example, Aznar-Lopez et al obtained 0.7% (10/1488) of positivity using the nested DimLis PCR RT-PCR method in oropharyngeal samples collected from five of the 27 bat species sampled throughout Spain between 2004 and 2010 (Aznar-Lopez et al., 2013). In this study, bat species found positive for rhabdoviruses were *Eptesicus isabellinus*, *Hypsugo savii*, *Miniopterus schreibersii*, *Plecotus auratus* and *Rhinolophus ferrumequinum*). Another study carried out in the *Desmodus rotundus* bat (vampire bat) in Guatemala using a generic hemi-nested PCR based on the primers PVO3/PVO4/ PVOnstF provided a positivity rate of 0.25% (1/396) after screening different samples (blood clot, serum, fecal or oral swab and urine) (Wray et al., 2016).

After Sanger sequencing, all the 23 positive samples exhibited the presence of a same rhabdovirus isolate, named Mediterranean bat virus (MBV) and belonging to the genus *Vesiculovirus*. The nearly complete genome sequence was retrieved for 16 samples (1 from Spain, 5 from Morocco and 10 from Algeria), and was characterized by a classical vesiculovirus organization with the presence of the five canonical genes (N, P, M, G, L). For each of these proteins, amino-acid identities were high (over 99%), demonstrating the close genetic relationships between these isolates. At the level of the genus *Vesiculovirus*, the genetic comparison between the other members revealed that Mediterranean bat virus can be considered as a new species. Indeed, MBV exhibited amino acid divergences of 33 %, 24.5 % and 37.6 % for the L, N and G proteins, respectively, with the most closely related vesiculovirus species *Vesiculovirus yinshui*. These genetic divergence results were in accordance with the ICTV demarcation criteria (minimum amino acid sequence divergence of 20%, 10% and 15% in L, N and G proteins, respectively) to validate the creation of a new virus species, and the *Vesiculovirus mediterranean* species was ratified in March 2022 (Kuhn, et al., 2022).

At the level of the genus *Vesiculovirus*, three main phylogroups were observed after phylogenetic analysis, among which all bat-related vesiculoviruses (including MBV isolates) clustered into the phylogroup III. Within this phylogroup, a clear geographical and/or bat species delimitation was observed between the origin of the different virus isolates, with one group gatherings together the American bat vesiculovirus (ABV) collected from *Eptesicus fuscus* in North America (Ng et al., 2013). Another cluster was composed of Chinese bat-related vesiculoviruses with Yinshui bat virus (YSBV) from *Rhinolophus sinicus*, and with Qiongzhong bat virus (QZBV) or Jinghong bat virus (JHBV), both obtained from *Rhinolophus affinis* (Luo et al., 2021a; Xu et al., 2018). The last one encompassed the different MBV isolates from the Mediterranean basin, collected from *Miniopterus schreibersii*, *Rhinolophus ferrumequinum* and *Rhinolophus euryale*. These results demonstrated that specific bat-related vesiculoviruses are actively circulating among various bat species worldwide.

The phylogenetic analysis of the different genome sequences of MBV isolates also revealed a geographical demarcation, with a distinct cluster of Algerian isolates, and a second cluster encompassing both the single Spanish strain and the isolates from Morocco. Among this latter cluster, the Spanish isolate diverged from the Moroccan ones, suggesting that the Spanish strain may have originated from Moroccan strains. Looking more closely at the scale of the caves, this demarcation was even more striking, since all the isolates in a single cave were distinct from those in other caves, even within the same country.

Unlike the other bat vesiculoviruses, MBV is not restricted to one bat species, being able to infect at least three ones so far, especially those of the genus *Rhinolophus*. All these three positive bat species are commonly found sharing the same roost. As an example for Morocco, caves III and IV host different mono-species bat colonies, with *Miniopterus schreibersii*, *Myotis emarginatus*, *Myotis punicus*, *Rhinolophus ferrumequinum* and *Rhinolophus euryale*. However, in these roosts, these species have often been observed in close contact, which may facilitate the diffusion and the circulation of MBV between individuals of different species. Furthermore, *Miniopterus schreibersii* is the only species capable of relatively long seasonal movements, sometimes covering distances in excess of 300 kilometers. It migrates between winter and summer roosts, potentially contributing to the dispersal of MBV in northern Africa, although little is known about its migration patterns in this region.

The migration of *Miniopterus schreibersii* is most documented in the Catalonia region of Spain, where Cave I is located. This roost hosts a large wintering bat colony consisting of 17,000 *Miniopterus schreibersii* and 150 *Rhinolophus ferrumequinum*. Towards the end of winter, *Miniopterus schreibersii* migrate to spring roosts and then to summer sites (Serra-Cobo et al., 1998). Thus, the bats in Cave I disperse over a vast territory, encompassing a significant part of Catalonia and southeastern France, potentially contributing to the dispersal and maintenance of MBV circulation. In addition, studies of other rhabdoviruses have demonstrated the importance of *Miniopterus schreibersii* in maintaining lyssavirus in colonies of other bat species (Pons-Salort et al., 2014; Colombi et al., 2019). It is possible that *Miniopterus schreibersii* plays a similar role with MBV, but further research is needed to confirm this hypothesis. Seroprevalence determination is a useful parameter to determine the level of bat exposure, and to try model the dynamics of circulation of this virus in populations, especially when the positivity rate of virus detection remains low. Indeed, such approaches have been conducted with success for other bat-borne viruses such as lyssavirus (i.e. *Lyssavirus hamburg*, EBLV-1) (Serra-Cobo et al., 2018).

Previous studies demonstrated that bat vesiculoviruses could be detected in different organs of infected individuals. For example, American bat vesiculovirus was identified in heart and lung homogenates, as well as in viscera pools of big brown bats (*Eptesicus fuscus*) (Ng et al., 2013). Chinese bat vesiculoviruses Jinghong bat virus and Benxi bat virus were detected in intestine and/or lung (Xu et al., 2018). Similarly, vesiculoviruses Yinshui bat virus, Taiyi bat virus and Qiongzhong bat virus, all identified in Chinese *Rhinolophus* bat species, were also detected in different organs (including, depending on the virus of interest, brain, heart, liver, spleen, lung, kidney and intestine) (Luo et al., 2021a).

In our study, we did not have access to any cadavers or organs. However, MBV was detected in two different types of biological samples, with saliva swabs for *Miniopterus schreibersii*, and with blood pellet for both *Rhinolophus euryale* and *Rhinolophus ferrumequinum*. Although limited, these data evidence that MBV can be excreted via saliva, at least in *Miniopterus* species, which could play a role in viral transmission. Unfortunately, the salivary swabs were placed directly into TRIzol after collection, which made it impossible to test this hypothesis using viral isolation tests.

The additional detection of MBV in blood pellet for *Rhinolophus euryale* and *Rhinolophus ferrumequinum* strongly suggest a viremia stage in both species. Most of vesiculovirus species was demonstrated to be transmitted by arthropods, mainly insects, which feed on the blood of infected vertebrates. Some studies have also demonstrated that these ectoparasites can host numerous viruses, including rhabdoviruses. For example, short rhabdoviral sequences were detected in different ectoparasites (*Nycteribia kolenatii*, *Nycteribia schmidli* and *Penicillidia conspicua*) collected on Spanish bats (Aznar-Lopez et al., 2013). In Uganda, two novel ledanteviruses, Bundibugyo and Kanyawara viruses, were identified in nycteribiid bat flies infesting pteropodid bats (Goldberg et al., 2017; Bennett et al., 2020). Kanyawara virus was also detected directly in one of these infested bats. Thus, bat flies or other arthropods may serve as ectoparasitic reservoirs of “bat-associated” viruses, which can play a direct role in the virus circulation in bat (similar to that observed for most vesiculoviruses and other mammals), or only transiently or sporadically infect bats (as appears to be the case with Kanyawara virus (Bennett et al., 2020). In our case, this could explain why MBV is found in different bat species sharing the same habitats. So far, the presence of the MBV in bat flies (*Nycteribiidae*) or ticks has not been investigated. In addition, virus isolation attempts using the remaining blood pellet, when still available, were not successful. This data will help to decipher the epidemiological cycle of MBV.

Various factors may explain the failure of attempts to isolate viruses from blood. The quality of the samples may be one of them, particularly in the case of blood tube bottoms from a retrospective collection, which may have undergone different freeze/thaw cycles. Moreover, the quantity available was very limited. New prospective samples would be required to ensure optimal conditions for success. In addition, alternative strategies need to be pursued, including the use of more appropriate substrates such as cell lines from the species concerned (*i.e. Miniopterus schreibersii*, *Rhinolophus euryale* and *Rhinolophus ferrumequinum*). In fact, MBV cell receptors are not known and could be more or less specific to different bat species. or insect cell lines. Similarly, arthropod cell lines could be used for virus isolation, although the potential arthropod vectors remain unknown.

In any case, the presence of MBV in the different positive individuals was not associated to any clinical manifestation or symptoms, these animals having been captured in flight using nets, then released after sampling. Another recent study also described the presence of vesiculovirus infection in Chinese bats without any clinical signs at the time of capture (Luo et al., 2021a). More generally, various studies have demonstrated virus infection in asymptomatic bats, underlying the fact that these animals are playing a major role as potential zoonotic reservoirs (Calisher et al., 2006; Letko et al., 2020; Van Brussel and Holmes, 2022). The current hypothesis is that a balanced immune response would enable them to maintain homeostasis during infection, limiting viral replication while avoiding the impact of excessive inflammation (Perrot and Dacheux, 2023; Irving et al. 2021). Deciphering these mechanisms, using adapted in vitro models, will help assess and avoid the potential zoonotic risk of these animals, while paving the way for the development of therapeutics for infectious and inflammatory diseases.

In conclusion, using a broad-spectrum detection technique, our study demonstrated that the prevalence of rhabdovirus within the bat samples tested remained low. Despite this low prevalence, we were able to demonstrate the presence of a new species among the genus *Vesiculovirus*, which is associated to three bat species with *Miniopterus schreibersii*, *Rhinolophus euryale* and *Rhinolophus ferrumequinum*. These data confirm that a specific group of vesiculoviruses circulates widely throughout the world in insectivorous bats, without being apparently associated with disease symptoms in these animals. According to the virus genus considered (*Vesiculovirus*) and to the nature of the samples found positive (particularly blood pellet), it is very likely that MBV can be considered as an arbovirus, like most other vesiculoviruses. This virus appears to be able to infect different species of bats, but its ability to infect other species within bats, or other mammals including humans, remains to be determined. This is especially true since the presence of this virus has been observed in saliva, suggesting that it could be excreted and transmitted via this mode. This potential zoonotic risk therefore requires more in-depth investigations, as do the mechanisms of infection and circulation in bat colonies. To achieve this goal, complementary analysis tools are in development, such as specific serological approaches, dedicated cellular models and reverse genetics approaches. These results also underline the role of bats are in rhabdovirus diffusion and highlight the importance to perform active surveillance of this animal reservoir, source of viral genetic diversity and potential rhabdovirus emergences.

## 5 Data availability

Sequences of the Mediterranean bat virus are publicly available in GenBank (accession numbers: MW491754-MW491760).

## 6 Supplementary data

Supplementary data are available at in the online site of the journal.

## Acknowledgements

We wish to thank the Natural Park of Sant Llorenç del Munt i l’Obac (Barcelona, Spain), the National Park of Chréa (Algeria), the National Park of Béjaia (Algeria) and Drs. Mehdi Elharrak, Bachir Harif, Sehhar El Ayachi, Elbia Abdelatif and Ahmim Mourad for their precious the contribution during field work. We would also like to thank all the other people who took part in the sample collection work in the field, without whom this study would not have been possible. We are also grateful to the Laboratory for Urgent Response to Biological Threats (CIBU) at Institut Pasteur, and in particular Christophe Batéjat, for providing us with different materials for the validation of the pan-rhabdo RT-nqPCR. Lastly, we thank Gaston for his excellent technical experience in bat design.

