## Supplemental material for "Large circulation of a novel vesiculovirus in bats in the Mediterranean region"

*** Correspondence:**

Dongsheng Luo

### 1 Supplementary Figures

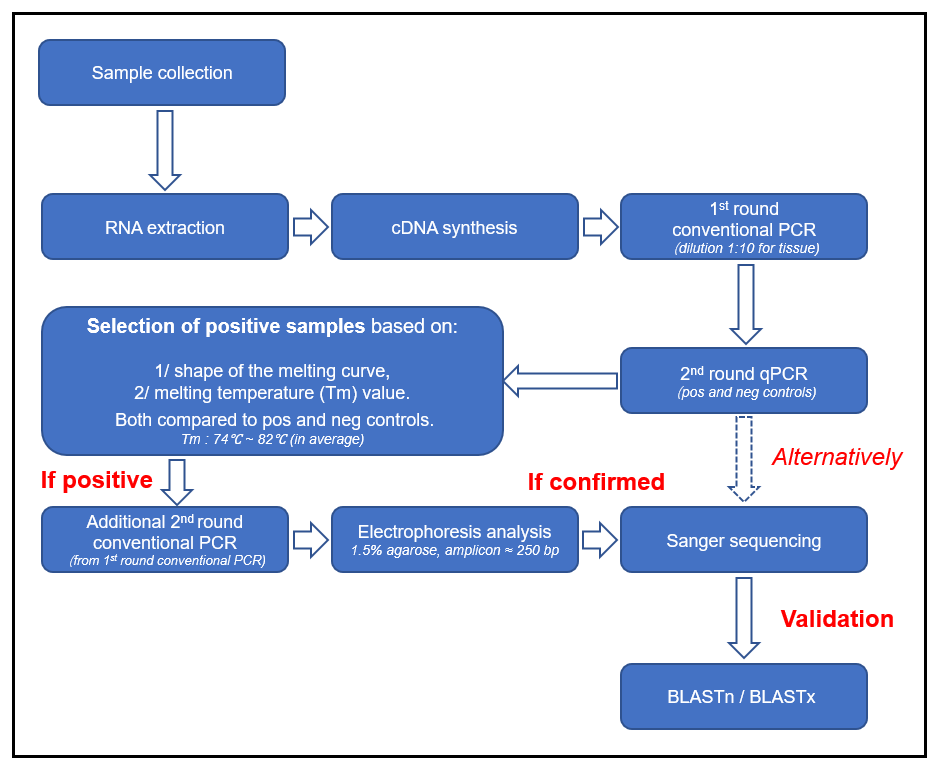

**Supplementary Figure 1.** Workflow analysis for the interpretation of the results obtained with the pan-rhabdo RT-nqPCR. After RNA extraction of the samples, a first cDNA step is performed, followed by a 1^st^ conventional PCR. A nested qPCR is systematically performed, and results are interpreted according to positive and negative samples, based on the shape of the melting curve and the value of the Tm. Positive samples are confirmed with an additional 2^nd^ round conventional PCR (from the 1^st^ round conventional PCR). Amplicons are analyzed after gel electrophoresis migration, and positive samples are Sanger sequenced. Validation of positive results is obtained after BLASTn and/or BLASTx analysis. Alternatively, amplicons from 2^nd^ round qPCR can also be directly Sanger sequenced and analyzed by BLAST. Templates (cDNA) from tissue samples need to be diluted 1:10 in nuclease-free water) before the 1^st^ round of conventional PCR. This dilution step is not required for liquid samples (e.g. blood or saliva).

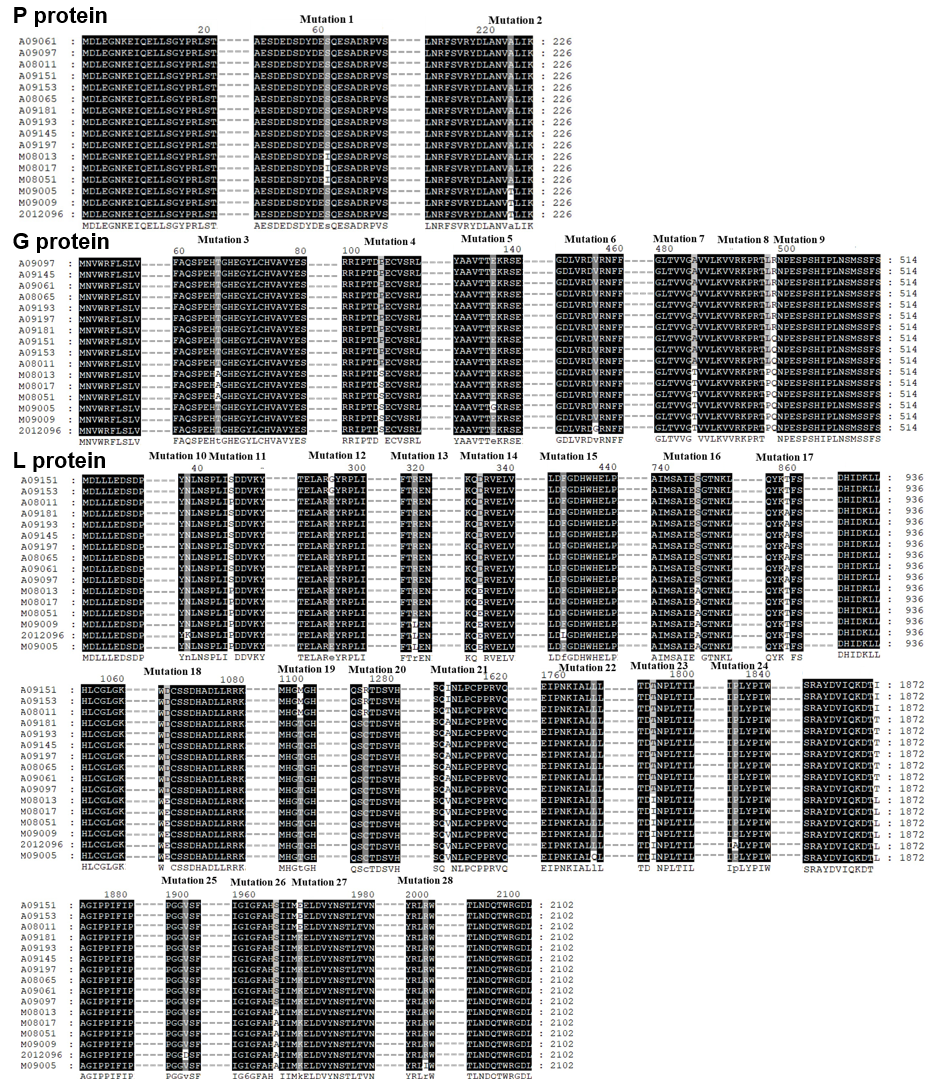

**Supplementary Figure 2.** Identification of the different deducted amino acid mutations between the 16 isolates of Mediterranean bat virus (MBV). Dot lines indicate the conserved regions not shown in the figure. A total of 28 amino acid mutations were identified and represented. The multiple alignment was performed with ClustalW 2.0).

### 2 Supplementary Tables

**Table S1.** Details of the bat samples included in this study, according to the species and to the type of samples (brain, saliva and blood).

| **Bat** | |  | **Sample type (No.)** | | | **Total (No.)** |
| --- | --- | --- | --- | --- | --- | --- |
| **Family** | **Species** |  | **Brain** | **Saliva** | **Blood** |  |
| Pteropodidae | *Rousettus aegyptiacus* |  |  |  | 100 | 100 |
|  | *Eidolon helvum* |  | 222 |  |  | 222 |
|  | *Epomophorus gambianus* |  | 5 | 7 |  | 12 |
|  | *Epomops buettikoferi* |  | 3 | 2 |  | 5 |
|  | *Epomops franqueti* |  | 3 |  |  | 3 |
|  | *Epomops sp.* |  | 1 | 4 |  | ~~5~~ |
|  | *Hypsignathus gambianus* |  | 1 |  |  | 1 |
|  | *Hypsignathus monstrosus* |  | 5 |  |  | 5 |
|  | *Megaloglossus azagnyi* |  | 1 |  |  | 1 |
|  | *Micropteropus pusilillus* |  | 99 | 41 |  | 140 |
|  | *Myonycteris leptodon* |  | 4 | 3 |  | 7 |
|  | *Myonycteris torquata* |  | 1 | 3 |  | 4 |
|  | *Nanonycteris veldkampii* |  | 3 | 1 |  | 4 |
|  | *Nanonycteris velkampi* |  | 1 |  |  | 1 |
|  | *Scotonycteris zenkeri* |  | 4 |  |  | 4 |
| Rhinolophidae | *Rhinolophus alcyone* |  | 3 | 2 |  | 5 |
|  | *Rhinolophus blasii* |  |  |  | 1 | 1 |
|  | *Rhinolophus euryale* |  |  |  | 41 | 41 |
|  | *Rhinolophus ferrumequinum* |  | 11 |  | 115 | 126 |
|  | *Rhinolophus hipposideros* |  |  |  | 1 | 1 |
|  | *Rhinolophus sp.* |  | 2 | 1 | 6 | 9 |
| Hipposideridae | *Hipposideros caffer* |  | 27 | 6 |  | 33 |
|  | *Hipposideros ruber* |  | 3 | 1 |  | 4 |
|  | *Hipposideros sp.* |  | 2 | 1 |  | 3 |
| Rhinopomatidae | *Rhinopoma hardwickii* |  |  |  | 7 | 7 |
|  | *Rhinopoma microphyllum* |  |  |  | 21 | 21 |
| Miniopteridae | *Miniopterus schreibersii* |  | 4 | 193 | 157 | 354 |
| Vespertilionidae | *Myotis blythii* |  |  | 1 | 12 | 13 |
|  | *Myotis capaccinii* |  |  | 7 | 7 | 14 |
|  | *Myotis dasycneme* |  | 1 |  |  | 1 |
|  | *Myotis emarginatus* |  | 3 |  | 27 | 30 |
|  | *Myotis escalerai* |  |  | 9 | 1 | 10 |
|  | *Myotis myotis* |  | 188 | 131 |  | 319 |
|  | *Myotis mystacinus* |  | 12 |  |  | 12 |
|  | *Myotis nattereri* |  | 1 |  |  | 1 |
|  | *Myotis punicus* |  | 1 |  | 297 | 298 |
|  | *Neoromicia nana* |  | 2 | 2 |  | 4 |
|  | *Noctulas noctula* |  | 48 |  |  | 48 |
|  | *Nyctalus leisleri* |  | 1 |  |  | 1 |
|  | *Nycteris grandis* |  | 1 | 1 |  | 2 |
|  | *Nycteris hispida* |  | 1 | 1 |  | 2 |
|  | *Nycteris thebaica* |  | 2 | 2 |  | 4 |
|  | *Nycteris sp.* |  | 2 |  |  | 2 |
|  | *Pipistrellus kuhlii* |  | 5 |  | 2 | 7 |
|  | *Pipistrellus pipistrellus* |  | 7 |  |  | 7 |
|  | *Scotophilus leucogaster* |  | 4 | 4 |  | 8 |
|  | *Vespertilio murinus* |  | 5 |  |  | 5 |
|  | *Eptesicus isabellinus* |  |  |  | 3 | 3 |
|  | *Eptesicus serotinus* |  | 32 |  |  | 32 |
| Emballonuridae | *Taphozous nudiventris* |  |  |  | 18 | 18 |
| Molossidae | *Molossus molossus* |  | 2 |  |  | 2 |
| **Total** | |  | 723 | 423 | 816 | 1962 |

**Table S2**. Description of the polymerase sequences from animal rhabdoviruses selected to design the primers of the pan-rhabdo RT-nqPCR.

| **Virus** | **Species** | **Genus** | **Location of first isolation** | **Year of isolation** | **Source species** | **GenBank accession** |
| --- | --- | --- | --- | --- | --- | --- |
| Arboretum virus (ABTV) | *Almendravirus arboretum* |  *Almendravirus* | Peru |  2009 | mosquitoe | NC_025393 |
| Puerto Almendras virus (PTAMV) | *Almendravirus almendras* | *Almendravirus* | Peru | 2009 | mosquitoe | NC_025395 |
| Curionopolis virus (CURV) | *Curiovirus curionopolis* |  *Curiovirus* | Brazil |  1985 | midge | NC_025354 |
| Iriri virus (IRIRV) | *Curiovirus iriri* |  *Curiovirus* | Brasil |  1982 | sandfly | NC_034544 |
| Itacaiunas virus (ITAV) | *Curiovirus itacaiunas* |  *Curiovirus* | Brazil |  1984 | midge | NC_034536 |
| Rochambeau virus (RBUV) | *Curiovirus rochambeau* |  *Curiovirus* | French Guiana | 1973 |  mosquitoe |  NC_034534 |
|  Adelaide River virus (ARV) |  *Ephemerovirus adelaide* |  *Ephemerovirus* | Australia |  1981 | bovine | NC_028246 |
| Berrimah virus (BRMV) | *Ephemerovirus berrimah* | *Ephemerovirus* | Australia | 1981 | bovine | NC_025358 |
| Bovine ephemeral fever virus (BEFV) | *Ephemerovirus febris* | *Ephemerovirus* | China | 2002 | bovine | KY315724 |
| Kimberley virus (KIMV) | *Ephemerovirus kimberley* | *Ephemerovirus* | Australia | 1980 | bovine | NC_025396 |
| Koolpinyah virus (KOOLV) | *Ephemerovirus koolpinyah* |  *Ephemerovirus* | Australia |  1985 | bovine | NC_028239 |
| Kotonkan virus (KOTV) | *Ephemerovirus kotonkan* | *Ephemerovirus* | Nigeria | 1967 | midge | NC_017714 |
| Malakal virus (MALV) | *Ephemerovirus kimberley* | *Ephemerovirus* | Sudan | 1963 | mosquitoe | JQ941707 |
| New Kent County virus (NKCV) | *Ephemerovirus kent* | *Ephemerovirus* | USA | 2016 | tick | MF615270 |
| Obodhiang virus (OBOV) | *Ephemerovirus obodhiang* | *Ephemerovirus* | Sudan | 1963 | mosquitoe | NC_017685 |
| Yata virus (YATV) | *Ephemerovirus yata* |  *Ephemerovirus* | Central African Republic |  1969 | mosquitoe | NC_028241 |
| Manitoba virus (MANV) | *Hapavirus manitoba* |  *Hapavirus* | Canada |  1977 | mosquitoe | NC_034531 |
| Ngaingan virus (NGAV) | *Hapavirus ngaingan* |  *Hapavirus* | Australia |  1970 | midge | NC_013955 |
| Gray Lodge virus (GLOV) | *Hapavirus graylodge* |  *Hapavirus* | USA |  1971 | mosquitoe | NC_034541 |
| Hart Park virus (HPV) | *Hapavirus hartpark* | *Hapavirus* | USA | 1955 | mosquitoe | NC_034447 |
| Joinjakaka virus (JOIV) | *Hapavirus joinjakaka* | *Hapavirus* | Papua-New Guinea | 1966 | mosquitoes | NC_034538 |
| Kamese virus (KAMV) | *Hapavirus kamese* | *Hapavirus* | Central African Republic | 1977 | mosquitoe | KX497133 |
| La Joya virus (LJV) | *Hapavirus lajoya* |  *Hapavirus* | Panama |  1958 | mosquitoe | NC_034537 |
| Landjia virus (LJAV) | *Hapavirus landjia* |  *Hapavirus* | Central African Republic |  1970 | bird | NC_034533 |
| Marco virus (MCOV) | *Hapavirus marco* |  *Hapavirus* | Brazil |  1962 | lizard | NC_034530 |
| Mosqueiro virus (MQOV) | *Hapavirus mosqueiro* | *Hapavirus* | Brazil | 1970 | mosquitoe | NC_034448 |
| Mossuril virus (MOSV) | *Hapavirus mossuril* |  *Hapavirus* | Mozambique |  1959 | mosquitoe | KM204993 |
| Ord River virus (ORV) | *Hapavirus ord* | *Hapavirus* | Australia | 1976 | mosquitoe | KY421920 |
| Parry Creek virus (PCV) | *Hapavirus parry* | *Hapavirus* | Australia | 1973 | mosquitoe | NC_034449 |
| Wongabel virus (WONV) | *Hapavirus wongabel* | *Hapavirus* | Australia | 1979 | midge | NC_011639 |
| Kolente virus (KOLEV) | *Ledantevirus kolente* | *Ledantevirus* | Guinea | 1985 | tick | NC_025342 |
| Kumasi rhabdovirus (KRV) | *Ledantevirus kumasi* | *Ledantevirus* | Ghana | 2011 | bat (*Eidolon helvum*) | NC_028236 |
| Le Dantec virus (LDV) | *Ledantevirus ledantec* |  *Ledantevirus* | Senegal |  1965 | human | NC_034443 |
| Nkolbisson virus (NKOV) | *Ledantevirus nkolbisson* | *Ledantevirus* | Cameroon | 1965 | mosquitoe | NC_034539 |
| Barur virus (BARV) | *Ledantevirus barur* |  *Ledantevirus* | India |  1962 | rat | NC_034535 |
| Fikirini virus (FKRV) | *Ledantevirus fikirini* |  *Ledantevirus* | Kenya |  2011 | bat (*Hipposideros commersoni*) | NC_025341 |
| Fukuoka virus (FUKV) | *Ledantevirus fukuoka* |  *Ledantevirus* | Japan |  1982 | midge | NC_034454 |
| Kern Canyon virus (KCV) | *Ledantevirus kern* | *Ledantevirus* | USA | 1968 | bat (*Myotis yumanensis*) | NC_034451 |
| Keuraliba virus (KEUV) | *Keuraliba virus* | *Ledantevirus* | Senegal | 1968 | gerbil | NC_034540 |
|  Mount Elgon bat virus (MEBV) |  *Ledantevirus elgon* |  *Ledantevirus* | Kenya |  1964 |  bat (*Rhinolophus hilderbrandtii eloquens*) |  NC_034545 |
| Nishimuro virus (NISV) | *Ledantevirus nishimuro* | *Ledantevirus* | Japan | 1962 | wild boar | AB609604 |
| Oita virus (OITAV) | *Ledantevirus oita* |  *Ledantevirus* | Japan |  1972 | bat (*Rhinolophus cornutus*) | NC_034548 |
| Wuhan louse fly virus 5 (WLFV5) | *Ledantevirus wuhan* | *Ledantevirus* | China | 2013 | fly | NC_031301 |
|  Yongjia tick virus 2 (YTV2) |  *Ledantevirus yongjia* | *Ledantevirus* | China | 2012 | tick | NC_031305 |
| Taiwan bat lyssavirus (TWBLV) | *Lyssavirus formosa* | *Lyssavirus* | Taiwan | 2016 | bat (*Pipistrellus abramus)* | MF472710 |
| Aravan virus (ARAV) | *Lyssavirus aravan* | *Lyssavirus* | Kyrgyzstan | 1991 | bat (*Myotis blythii*) | NC_020808 |
| Australian bat lyssavirus (ABLV) | *Lyssavirus australis* | *Lyssavirus* | Australia | 1996 | bat (*Saccolaimus flaviventris*) | NC_003243 |
| Bokeloh bat lyssavirus (BBLV) | *Lyssavirus bokeloh* | *Lyssavirus* | Germany | 2010 | bat (*Myotis nattererii*) | NC_025251 |
| Duvenhage virus (DUVV) | *Lyssavirus duvenhage* | *Lyssavirus* | South Africa | 1971 | human | NC_020810 |
| European bat lyssavirus 1 (EBLV1) | *Lyssavirus hamburg* | *Lyssavirus* | Germany | 1968 | bat (*Eptesicus serotinus*) | EF157976 |
| European bat lyssavirus 2 (EBLV2) | *Lyssavirus helsinki* | *Lyssavirus* | United Kingdom | 2016 | bat (*Myotis daubentonii*) | KY688156 |
| Gannoruwa bat lyssavirus (GBLV) | *Lyssavirus gannoruwa* |  *Lyssavirus* | Sri Lanka |  2015 |  bat (*Pteropus giganteus*) |  NC_031988 |
| Ikoma lyssavirus (IKOV) | *Lyssavirus ikoma* | *Lyssavirus* | Tanzania | 2009 | African civet | NC_018629 |
| Irkut virus (IRKV) | *Lyssavirus irkut* | *Lyssavirus* | Russia | 2002 | bat (*Murina leucogaster*) | NC_020809 |
| Khujand virus (KHUV) | *Lyssavirus khujand* | *Lyssavirus* | Tajikstan | 2001 | bat (*Myotis mystacinus*) | NC_025385 |
| Lagos bat virus (LBV) | *Lyssavirus lagos* | *Lyssavirus* | Ghana | 2015 | bat (*Eidolon helvum*) | LN849915 |
| Lleida bat lyssavirus (LLEBV) | *Lyssavirus lleida* |  *Lyssavirus* | Spain | 2011 |  bat (*Miniopterus schreibersii*) |  NC_031955 |
| Mokola virus (MOKV) | *Lyssavirus mokola* |  *Lyssavirus* | South Africa |  1996 | feline | KF155008 |
| Rabies virus (RABV) | *Lyssavirus rabies* |  *Lyssavirus* | France |  1983 | human | EU293121 |
| Shimoni bat virus (SHIBV) | *Lyssavirus shimoni* |  *Lyssavirus* | Kenya |  2009 | bat (*Hipposideros commersoni*) | NC_025365 |
| West Caucasian bat virus (WCBV) | *Lyssavirus caucasicus* |  *Lyssavirus* | Russia | 2001 | bat (*Miniopterus schreibersii*) | NC_025377 |
| Moussa virus (MOUV) | *Mousrhavirus moussa* | *Mousrhavirus* | Cote d'Ivoire | 2004 | mosquitoe | NC_025359 |
| Eel virus European X (EVEX) | *Perhabdovirus anguilla* | *Perhabdovirus* | Denmark | 1986 | fish | KC608035 |
| Perch rhabdovirus (PRV) | *Perhabdovirus perca* | *Perhabdovirus* | France | 1981 | fish | NC_020803 |
| Drosophila affinis sigmavirus (DAffSV) | *Sigmavirus affinis* | *Sigmavirus* | USA | 2007 | fly | KR822811 |
| Drosophila melanogaster sigmavirus (DMelSV) | *Sigmavirus melanogaster* | *Sigmavirus* | USA | unknown | fly | JX403934 |
| Drosophila obscura sigmavirus (DObSV) | *Sigmavirus obscura* | *Sigmavirus* | United Kingdom | 2007 | fly | NC_022580 |
| Carp sprivivirus (CSV) | *Sprivivirus cyprinus* | *Sprivivirus* | China | 2016 | fish | KY475636 |
| Grass carp rhabdovirus (GrCRV) | *Sprivivirus esox* | *Sprivivirus* | Germany | 1982 | fish | NC_025376 |
| Pike fry rhabdovirus (PFRV) | *Sprivivirus esox* | *Sprivivirus* | France | 1972 | fish | NC_025356 |
| Tench rhabdovirus (TenRV) | *Sprivivirus esox* | *Sprivivirus* | Germany | 1982 | fish | NC_025371 |
| Sena Madureira virus (SMV) | *Sripuvirus madureira* | *Sripuvirus* | Brazil | 1976 | lizard | NC_034529 |
| Sripur virus (SRIV) | *Sripuvirus sripur* | *Sripuvirus* | India | 1973 | sandfly | NC_034542 |
| Almpiwar virus (ALMV) | *Sripuvirus almpiwar* | *Sripuvirus* | Australia | 1966 | lizard | NC_025391 |
| Chaco virus (CHOV) | *Sripuvirus chaco* | *Sripuvirus* | Brazil | 1962 | lizard | NC_034550 |
| Niakha virus (NIAV) | *Sripuvirus niakha* | *Sripuvirus* | Senegal | 1992 | sandfly | NC_025405 |
| Oak Vale virus (OVV) | *Sunrhavirus oakvale* | *Sunrhavirus* | Australia | 1982 | mosquitoe | NC_025399 |
| Ekpoma virus 2 (EKV2) | *Tibrovirus betaekpoma* | *Tibrovirus* | Nigeria | 2011 | human | KP324828 |
| Bas-Congo virus (BASV) | *Tibrovirus congo* | *Tibrovirus* | Democratic Republic of the Congo | 2009 | human | JX297815 |
| Coastal Plains virus (CPV) | *Tibrovirus coastal* | *Tibrovirus* | Australia | 1981 | bovine | NC_025397 |
| Ekpoma virus (EKV) | *Unclassified* | *Tibrovirus* | China | 2017 | human | MF079256 |
| Ekpoma virus 1 (EKV1) | *Tibrovirus alphaekpoma* | *Tibrovirus* | Nigeria | 2011 | human | KP324827 |
| Sweetwater Branch virus (SWBV) | *Tibrovirus coastal* | *Tibrovirus* | USA | 1982 | midge | NC_034546 |
| Tibrogargan virus (TIBV) | *Tibrovirus tibrogargan* | *Tibrovirus* | Australia | 1976 | midge | NC_020804 |
| Durham virus (DURV) | *Tupavirus durham* | *Tupavirus* | USA | 2005 | bird | FJ952155 |
| Klamath virus (KLAV) | *Tupavirus klamath* | *Tupavirus* | USA | 1962 | vole | NC_034549 |
| Tupaia virus (TUPV) | *Tupavirus tupaia* | *Tupavirus* | Thailand | unknown | tree shrew | NC_007020 |
| American bat vesiculovirus (ABVV) | *Vesiculovirus eptesicus* | *Vesiculovirus* | USA | 2008 | bat (*Eptesicus fuscus*) | NC_022755 |
| Carajas virus (CARV) | *Vesiculovirus carajas* | *Vesiculovirus* | Brazil | 1983 | sandfly | KM205015 |
| Chandipura virus (CHPV) | *Vesiculovirus chandipura* | *Vesiculovirus* | India | 2004 | human | NC_020805 |
| Cocal virus (COCV) | *Vesiculovirus cocal* | *Vesiculovirus* | Trinidad | 1961 | rodent | EU373657 |
| Isfahan virus (ISFV) | *Vesiculovirus isfahan* | *Vesiculovirus* | Iran | 1975 | sandfly | NC_020806 |
| Jurona virus (JURV) | *Vesiculovirus jurona* | *Vesiculovirus* | Brazil | 1962 | human | NC_025392 |
| Malpais Spring virus (MSPV) | *Vesiculovirus malpais* | *Vesiculovirus* | USA | 1985 | mosquitoe | NC_025364 |
| Maraba virus (MARAV) | *Vesiculovirus maraba* | *Vesiculovirus* | Brazil | 1983 | sandfly | NC_025255 |
| Morreton virus (MORV) | *Vesiculovirus morreton* | *Vesiculovirus* | Colombia | 1986 | sandfly | NC_034508 |
| Perinet virus (PERV) | *Vesiculovirus perinet* | *Vesiculovirus* | Madagascar | 2008 | mosquito | NC_025394 |
| Piry virus (PIRYV) | *Vesiculovirus piry* | *Vesiculovirus* | Brazil | 1960 | Philander opossum | KU178986 |
| Radi virus (RADV) | *Vesiculovirus radi* | *Vesiculovirus* | Italy | 1982 | sandfly | KM205024 |
| Vesicular stomatitis Alagoas virus (VSAV) | *Vesiculovirus indiana* | *Vesiculovirus* | USA | 1998 | horse | NC_001560 |
| Vesicular stomatitis New Jersey virus (VSNJV) | *Vesiculovirus newjersey* | *Vesiculovirus* | Honduras | 1984 | bovine | NC_024473 |
| VS Alagoas virus (VSAV) | *Vesiculovirus alagoas* | *Vesiculovirus* | Brazil | 1964 | mule | NC_025353 |
| Yug Bogdanovac virus (YBV) | *Vesiculovirus bogdanovac* | *Vesiculovirus* | Serbia | 2011 | Vero-E6 cells | NC_025378 |

**Table S3.** Description of the primers tested during the validation step of the different pan-rhabdovirus PCR systems.

| **PCR system** | **Type** | **Primer** | **Sequence (5'-3')** | **Length (nt)** | **Reference position** | **Amplicon length (bp)^b^** | **Reference** |
| --- | --- | --- | --- | --- | --- | --- | --- |
| Screen-rhabdo qPCR_1 | SYBR Green-qPCR | F1_Rhabdovirus | ATWGGNYTNAARSSIAARGA | 20 | 1633-1652^a^ | 155 | This study |
|  |  | 2000R1_ Rhabdovirus | ADRTYRTCNGYCATIGTIA | 19 | 1769-1787^a^ |  |  |
| Screen-rhabdo qPCR_2 | SYBR Green-qPCR | 2100F2_ Rhabdovirus | TNGAYTAYGANAARTGGAAIAA | 22 | 1871-1892^a^ | 121 | This study |
|  |  | 2220R2_ Rhabdovirus | TTYTBAAANAAITYRTGIG | 19 | 1973-1991^a^ |  |  |
| Screen-rhabdo qPCR_3 | SYBR Green-qPCR | 2340F3_ Rhabdovirus | GARGGNYTNMGDCARAAIGGITGG | 24 | 2098-2121^a^ | 119 | This study |
|  |  | R1_Rhabdovirus | RYYTGRTTRTCNCCYTGIGC | 20 | 2188-2207^a^ |  |  |
| Screen-rhabdo qPCR_4 | SYBR Green-qPCR | F2_Rhabdovirus-M | GAYTAYGANAARTGGAAYAAYYAYCA | 26 | 1873-1898^a^ | 239 | This study |
|  |  | R2_Rhabdovirus | TGYCKNARNCCYTCYARNCCICC | 23 | 2089-2111^a^ |  |  |
| Pan-rhabdo RT-nqPCR | Conventional PCR (Nest first round) | F1_Rhabdovirus | ATWGGNYTNAARSSIAARGA | 20 | 1633-1652^a^ | 575 | This study |
|  |  | R1_Rhabdovirus | RYYTGRTTRTCNCCYTGIGC | 20 | 2188-2207^a^ |  |  |
|  | SYBR Green-qPCR/ Conventional PCR (Nest second round) | F2_Rhabdovirus-M | GAYTAYGANAARTGGAAYAAYYAYCA | 26 | 1873-1898^a^ | 239 |  |
|  |  | R2_Rhabdovirus | TGYCKNARNCCYTCYARNCCICC | 23 | 2089-2111^a^ |  |  |
| Pan-rhabdo RT-nPCR_1 | Conventional PCR (Nest first round) | DimLis1F | GGKMGRTTYTTYKCHYTDATG | 21 | 1650–1670^b^ | 466 | Aznar-Lopez *et al.*, 2013 |
|  |  | DimLis1R | CARAARGGNTGGASYNTHBT | 20 | 2097-2116^b^ |  |  |
|  | Conventional PCR (Nest second round) | DimLis2F | YTNTTYVANGSVYTRACNATG | 21 | 1734–1754^b^ | 150 |  |
|  |  | DimLis2R | TGGAAYAAYCAYCARMGRHWD | 21 | 1854–1874^b^ |  |  |
| Pan-rhabdo RT-nPCR_2 | Conventional PCR (Nest first round) | PVO3 | CCADMCBTTTTGYCKYARRCCTTC | 24 | 1655–1678^b^ | 458 | Wray *et al.*, 2016 |
|  |  | PVO4 | RAAGGYAGRTTTTTYKCDYTRATG | 24 | 2090-2113^b^ |  |  |
|  | Conventional PCR (Nest second round) | PVO3 | CCADMCBTTTTGYCKYARRCCTTC | 24 | 1655–1678^b^ | 260 |  |
|  |  | PVOnstF | AARTGGAAYAAYCAYCARMG | 20 | 1896-1915^b^ |  |  |

R=A/G, Y=C/T, M=A/C, K=G/T, S=G/C, W=A/T, H=A/T/C, B=G/T/C, V=G/A/C, D=G/A/T, N=A/T/C/G and I (hypoxanthine).

^a^ According to the Drosophila obscura sigmavirus (DObSV) L gene nucleotide sequence (GenBank accession number NC022580).

^b^ According to the rabies virus (RABV, CVS strain) L gene nucleotide sequence (GenBank accession number GQ918139).

**Table S4.** Specific primers designed to complete the full-length genome sequences of the bat Mediterranean vesiculovirus (MBV) isolates.

| **Target sample** | **Primer** | **Sequence** | **Size (bp)** |
| --- | --- | --- | --- |
| 2012096^a^ | 2012096_N-terminal_F1 | 5' TTGTCCCAAAGTGTCTCGCG 3' | 560 |
|  | 2012096_N-terminal_R1 | 5' CATTAGAAGAGCCCGATACTCTGG 3' |  |
|  | 2012096_N-terminal_F2 | 5' TCTCGCGTCCCGACCACC 3' |  |
|  | 2012096_N-terminal_R2 | 5' GAGGAGGTATGTCGGTAGCC 3' |  |
|  | 2012096_M-G_F1 | 5' CCTGCAGACATAGTTAAGTGGGC 3' | 1300 |
|  | 2012096_M-G_R1 | 5' GTGTCCTCATACACTGTTTTACAGGG 3' |  |
|  | 2012096_M-G_F2 | 5' CAGATTCTCAGTCCGTTATGATCTGGC 3' |  |
|  | 2012096_M-G_R2 | 5' TCATCCACTCCCACGTGGTGTGGG 3' |  |
|  | 2012096_L-1_F1 | 5' GACTGGAAGGGCCCGGATGATC 3' | 1700 |
|  | 2012096_L-1_R1 | 5' TCTACCACCCTCATCAAATCAGG 3' |  |
|  | 2012096_L-1_F2 | 5' GCCTGGTTTGCATCATGGATTGG 3' |  |
|  | 2012096_L-1_R2 | 5' TGGGGTAACCTAAAAACTGGCC 3' |  |
|  | 2012096_L-2_F1 | 5' CCAAAATGAAGTCATACGTGATTC 3' | 1500 |
|  | 2012096_L-2_R1 | 5' CTGAGTGGACAAAACAAAAGGCC 3' |  |
|  | 2012096_L-2_F2 | 5' CCTATGGTCCATTGACCCCTTATTTCCC 3' |  |
|  | 2012096_L-2_R2 | 5' GCAGTGTCAGAAAATATCCAAATCGCAGG 3' |  |
|  | 2012096_L-3_F1 | 5' GGATGTGTACAACTCAACATTAACGG 3' | 500 |
|  | 2012096_L-3_R1 | 5' CCTGATTCATAAATCCCCCCTCC 3' |  |
|  | 2012096_L-3_F2 | 5' CGATTCAAAATATCGACTAAGATGG 3' |  |
|  | 2012096_L-3_R2 | 5' CCCCACAGAAGCAGATGGTACGG 3' |  |
| A09145^b^ | A09145_N-terminal_F1 | 5' TTGTCCCAAAGTGTCTCGCG 3' | 360 |
|  | A09145_N-terminal_R1 | 5' TGTCCATATTCTCCTCCGCG 3' |  |
|  | A09145_N-terminal_F2 | 5' TCCCGACCACCCATTGATTG 3' |  |
|  | A09145_N-terminal_R2 | 5' ATAGTTATGCCGAAGGATGTCC 3' |  |

^a^ Virus isolate from Spanish bat saliva sample.

^b^ Virus isolate from Algerian bat blood sample.

**Table S5.** List of vesiculoviruses and sequence identification selected for the phylogenetic analysis of Mediterraean bat vesiculovirus isolates. The lyssavirus RABV was selected as outlier.

| **Virus (acronym)** | **Genus** | **Location of first isolation** | **Year of isolation** | **Host species** | **GenBank accession** |
| --- | --- | --- | --- | --- | --- |
| Rabies lyssavirus (RABV) | *Lyssavirus* | France | 1983 | Human/Bat | EU293121 |
| Piry vesiculovirus(PIRV) | *Vesiculovirus* | Brazil | 1960 | Gray four-eyed opossums (*Philander opossum*) | KU178986 |
| Jurona vesiculovirus (JURV) | *Vesiculovirus* | Brazil | 1962 | Human | NC025392 |
| Carajas vesiculovirus (CARV) | *Vesiculovirus* | Brazil | 1983 | Sandfly (*Lutzomyia sp.*) | KM205015 |
| Maraba vesiculovirus (MARAV) | *Vesiculovirus* | Brazil | 1983 | Sandfly | NC025255 |
| VS Alagoas vesiculovirus (VSAV) | *Vesiculovirus* | Brazil | 1964 | Mule | NC025353 |
| Jinghong bat vesiculovirus (JHBV) | *Vesiculovirus* | China | 2011 | Bat (*Rhinolophus affinis*) | MF279192 |
| Morreton vesiculovirus (MORV) | *Vesiculovirus* | Colombia | 1986 | Sandfly (*Lutzomyia sp.*) | NC034508 |
| VS New Jersey vesiculovirus (VSNJV) | *Vesiculovirus* | Honduras | 1984 | Bovine | NC024473 |
| Chandipura vesiculovirus (CHNV) | *Vesiculovirus* | India | 2004 | Human | NC020805 |
| Isfahan vesiculovirus (ISFV) | *Vesiculovirus* | Iran | 1975 | Sandfly (*Phlebotomus papatasi*) | NC020806 |
| Radi vesiculovirus (RADIV) | *Vesiculovirus* | Italy | 1982 | Sandfly (*Phlebotomus perfiliewi*) | KM205024 |
| Perinet vesiculovirus (PERV) | *Vesiculovirus* | Madagascar | 2008 | Mosquito | NC025394 |
| Yug Bogdanovac vesiculovirus (YBV) | *Vesiculovirus* | Serbia | 2011 | Sandfly (Vero-E6 cells) | NC025378 |
| Cocal vesiculovirus (COCV) | *Vesiculovirus* | Trinidad | 1961 | Rodent | EU373657 |
| American bat vesiculovirus (ABV) | *Vesiculovirus* | USA | 2008 | Bat (*Eptesicus fuscus*) | NC022755 |
| Malpais Spring vesiculovirus (MSPV) | *Vesiculovirus* | USA | 1985 | Mosquito (*Aedes campestris*) | NC025364 |
| VS Indiana vesiculovirus (VSIV) | *Vesiculovirus* | USA | 1998 | Horse | NC001560 |
| Qiongzhong bat virus 1127 (QZBV_1127) | *Vesiculovirus* | China | 2007 | Bat (*Rhinolophus affinis*) | MN607593 |
| Yinshui bat virus 1017 (YSBV_1017) | *Vesiculovirus* | China | 2007 | Bat (*Rhinolophus sinica*) | MN607594 |
| Yinshui bat virus D170001 (YSBV_D170001) | *Vesiculovirus* | China | 2018 | Bat (*Rhinolophus sinica*) | MN607595 |
| Yinshui bat virus D170022 (YSBV_D170022) | *Vesiculovirus* | China | 2019 | Bat (*Rhinolophus sinica*) | MN607596 |
| Yinshui bat virus D170190 (YSBV_D170190) | *Vesiculovirus* | China | 2020 | Bat (*Rhinolophus sinica*) | MN607597 |
| Bughendera virus (BUGV) | *Ledantevirus* | Uganda | 2017 | Batfly (Dipseliopoda sp.) | MT325641 |
| Mediterranean bat virus (MBV_2012096) | *Vesiculovirus* | Spain | 2012 | Bat (*Miniopterus schreibersii*) | MW557328 (this studiy) |
| Mediterranean bat virus (MBV_A08011) | *Vesiculovirus* | Algeria | 2008 | Bat (*Rhinolophus ferrumequinum*) | MW557329 (this study) |
| Mediterranean bat virus (MBV_A08065) | *Vesiculovirus* | Algeria | 2008 | Bat (*Rhinolophus euryale*) | MW557330 (this study) |
| Mediterranean bat virus (MBV_A09061) | *Vesiculovirus* | Algeria | 2009 | Bat (*Rhinolophus ferrumequinum*) | MW557331 (this study) |
| Mediterranean bat virus (MBV_A09097) | *Vesiculovirus* | Algeria | 2009 | Bat (*Rhinolophus ferrumequinum*) | MW557332 (this study) |
| Mediterranean bat virus (MBV_A09145) | *Vesiculovirus* | Algeria | 2009 | Bat (*Rhinolophus ferrumequinum*) | MW557333 (this study) |
| Mediterranean bat virus (MBV_A09151) | *Vesiculovirus* | Algeria | 2009 | Bat (*Rhinolophus ferrumequinum*) | MW557334 (this study) |
| Mediterranean bat virus (MBV_A09153) | *Vesiculovirus* | Algeria | 2009 | Bat (*Rhinolophus ferrumequinum*) | MW557335 (this study) |
| Mediterranean bat virus (MBV_A09181) | *Vesiculovirus* | Algeria | 2009 | Bat (*Rhinolophus ferrumequinum*) | NC_076939 (this study) |
| Mediterranean bat virus (MBV_A09193) | *Vesiculovirus* | Algeria | 2009 | Bat (*Rhinolophus ferrumequinum*) | MW557337 (this study) |
| Mediterranean bat virus (MBV_A09197) | *Vesiculovirus* | Algeria | 2009 | Bat (*Rhinolophus ferrumequinum*) | MW557338 (this study) |
| Mediterranean bat virus (MBV_M08013) | *Vesiculovirus* | Morocco | 2008 | Bat (*Rhinolophus ferrumequinum*) | MW557339 (this study) |
| Mediterranean bat virus (MBV_M08017) | *Vesiculovirus* | Morocco | 2008 | Bat (*Rhinolophus ferrumequinum*) | MW557340 (this study) |
| Mediterranean bat virus (MBV_M08051) | *Vesiculovirus* | Morocco | 2008 | Bat (*Rhinolophus ferrumequinum*) | MW557341 (this study) |
| Mediterranean bat virus (MBV_M09005) | *Vesiculovirus* | Morocco | 2009 | Bat (*Rhinolophus ferrumequinum*) | MW557342 (this study) |
| Mediterranean bat virus (MBV_M09009) | *Vesiculovirus* | Morocco | 2009 | Bat (*Rhinolophus ferrumequinum*) | MW557343 (this study) |
| American bat vesiculovirus strain 20-0654 (ABV_20-0654) | *Vesiculovirus* | USA | 2020 | Bat (*Eptesicus fuscus*) | MT561344 |
| American bat vesiculovirus strain 20-7011 (ABV_20-7011) | *Vesiculovirus* | USA | 2020 | Bat (*Eptesicus fuscus*) | MT561345 |
| American bat vesiculovirus strain 20-7177 (ABV_20-7177) | *Vesiculovirus* | USA | 2020 | Bat (*Eptesicus fuscus*) | MT561346 |
| American bat vesiculovirus strain 20-8862 (ABV_20-8862) | *Vesiculovirus* | USA | 2020 | Bat (*Eptesicus fuscus*) | MT561347 |

**Table S6.** Determination of the optimal concentration of primers for each pan-rhabdo PCR system under initial validation, using the reference strain Piry virus (PIRYV, BeAn 24232, 0413BRE) (*Vesiculovirus* genus). Tm values systems are indicated for each qPCR.

| **Primer concentration^a^** | **Sample dilution** | **PCR system** | | | | |
| --- | --- | --- | --- | --- | --- | --- |
|  |  | **Screen-rhabdo qPCR_1 (Tm=50°C)** | **Screen-rhabdo qPCR_2 (Tm=48°C)** | **Screen-rhabdo qPCR_3 (Tm=58°C)** | **Screen-rhabdo qPCR_4 (Tm=55°C)** | **Pan-rhabdo RT-nqPCR (1^st^ round, Tm=50°C) (2^nd^ round, Tm=56°C)** |
| 200 μM | 1 | Positive | Positive | Positive | Positive | Positive |
|  | 1:10 | Positive | Positive | Positive | Positive | Positive |
|  | 1:100 | Negative | Negative | Negative | Negative | Positive |
| 20 μM | 1 | Positive | Positive | Positive | Positive | Positive |
|  | 1:10 | Negative | Negative | Negative | Positive | Positive |
|  | 1:100 | Negative | Negative | Negative | Negative | Negative |

^a^ Final concentration of each primer (50 µL mix reaction).

**Table S7.** Description of the samples (rhabdoviruses and field samples) and the results obtained after the comparative evaluation of the different primers and qPCR systems.

| **Virus isolate** | **Strain** | **Species** | **Genus** | **Origin** | **Sample type** | **PCR results from different assay** | | | | | | |
| --- | --- | --- | --- | --- | --- | --- | --- | --- | --- | --- | --- | --- |
|  |  |  |  |  |  | **Rhabdo-screening qPCR_1** | **Rhabdo-screening qPCR_2** | **Rhabdo-screening qPCR_3** | **Rhabdo-screening qPCR_4** | **Pan-rhabdo-RT-nqPCR** | **Rhabdo-screening nest conventional PCR_1** | **Rhabdo-screening nest conventional PCR_2** |
| Sandjimba virus (SJAV) | DakAnB 373d (0408RCA) | *Sunrhavirus sandjimba* | *Sunrhavirus* | Bird (*Acrocephalus schoenbaeus*) | Brain (mouse)^c^ | Negative | Negative | Negative | Positive | Positive | ND | ND |
| Piry virus (PIRV) | BeAn 24232 (0413BRE) | *Vesiculovirus piry* | *Vesiculovirus* | Gray four-eyed opossums (*Philander opossum*) | Brain (mouse)^c^ | Positive | Positive | Positive | Positive | Positive | Negative | Positive |
| Jurona virus (JURV) | BeAr 40578 (0414BRE) | *Vesiculovirus jurona* | *Vesiculovirus* | Human | Brain (mouse)^c^ | Positive | Positive | Positive | Positive | Positive | ND | ND |
| Nkolbisson virus (NKOV) | Ar YM 31/65 (0425CAM) | *Ledantevirus nkolbisson* | *Ledantevirus* | Mosquito (*Eretmapodites leucopous*) | Brain (mouse)^c^ | Negative | Negative | Negative | Positive | Positive | ND | ND |
| Keuraliba virus (KEUV) | DakAnD 5314 (9715SEN) | *Ledantevirus keuraliba* | *Ledantevirus* | Gerbil (*Tatera kempi*) | Brain (mouse)^c^ | Negative | Positive | Positive | Positive | Positive | ND | ND |
| Bovine ephemeral fever virus (BEFV) | 7635MAY | *Ephemeroviru febris* | *Ephemerovirus* | Bovine | Blood^d^ | Negative | Negative | Negative | Negative | Positive | ND | ND |
| Bovine ephemeral fever virus (BEFV) | 7645MAY | *Ephemerovirus febris* | *Ephemerovirus* | Bovine | Blood^d^ | Negative | Positive | Negative | Positive | Positive | ND | ND |
| Vesicular stomatitis New Jersey virus (VSNJV) | VSV NJ-O (05004FRA) | *Vesiculovirus newjersey* | *Vesiculovirus* | Bovine | Brain (mouse)^c^ | ND^b^ | ND | ND | ND | Positive | Negative | Negative |
| Mediterranean bat virus (MBV) | A09061 | *Vesiculovirus mediterranean* | *Vesiculovirus* | Bat (*Rhinolophus ferrumequinum*) | Blood^d^ | ND | ND | ND | Negative | Positive | Negative | Positive |
| Mediterranean bat virus (MBV | A09097 | *Vesiculovirus mediterranean* | *Vesiculovirus* | Bat (*Rhinolophus ferrumequinum*) | Blood^d^ | ND | ND | ND | Negative | Positive | Negative | Positive |
| NA^a^ | E08157 | NA | NA | Bat (*Rousettus aegyptiacus*) | Blood^d^ | ND | ND | ND | ND | Negative | Negative | Negative |
| NA | E08159 | NA | NA | Bat (*Rousettus aegyptiacus*) | Blood^d^ | ND | ND | ND | ND | Negative | Negative | Negative |
| NA | E08195 | NA | NA | Bat (*Rousettus aegyptiacus*) | Blood^d^ | ND | ND | ND | ND | Negative | Negative | Negative |
| NA | A09027 | NA | NA | Bat (*Miniopterus schreibersii*) | Blood^d^ | ND | ND | ND | ND | Negative | Negative | Negative |
| NA | A09037 | NA | NA | Bat (*Miniopterus schreibersii*) | Blood^d^ | ND | ND | ND | ND | Negative | Negative | Negative |
| NA | A09039 | NA | NA | Bat (*Miniopterus schreibersii*) | Blood^d^ | ND | ND | ND | ND | Negative | Negative | Negative |
| NA | D0T039 | NA | NA | Bat (*Micropteropus pusillus* ) | Saliva^d^ | ND | ND | ND | ND | Negative | Negative | Negative |
| NA | DOT042 | NA | NA | Bat (*Micropteropus pusillus* ) | Saliva^d^ | ND | ND | ND | ND | Negative | Negative | Negative |
| NA | DOT061 | NA | NA | Bat (*Micropteropus pusillus* ) | Saliva^d^ | ND | ND | ND | ND | Negative | Negative | Negative |
| NA | KAB11 | NA | NA | Bat (*Micropteropus pusillus* ) | Saliva^d^ | ND | ND | ND | ND | Negative | Negative | Negative |

^a^ NA: Not applicable (negative sample).

^b^ ND: Not done.

^c^ Laboratory sample.

^d^ Field sample

**Table S8:** Details and NGS results of the 23 bat samples positive for rhabdovirus detection by the Pan-rhabdo RT-nqPCR.

| **Sample number** | **Sample type** | **Bat species** | **Sex** | **Country** | **Cave** | **Collection date** | **NGS result** |
| --- | --- | --- | --- | --- | --- | --- | --- |
| 2012086 | Saliva | *Miniopterus schreibersii* | Male | Spain | I (A. Daví) | 10/12/2012 | Few reads |
| 2012088 | Saliva | *Miniopterus schreibersii* | Female | Spain | I (A. Daví) | 10/12/2012 | Few reads |
| 2012094 | Saliva | *Miniopterus schreibersii* | Female | Spain | I (A. Daví) | 10/12/2012 | Few reads |
| 2012096 | Saliva | *Miniopterus schreibersii* | Male | Spain | I (A. Daví) | 10/12/2012 | Nearly full genome |
| 2012098 | Saliva | *Miniopterus schreibersii* | Male | Spain | I (A. Daví) | 10/12/2012 | Few reads |
| 2012100 | Saliva | *Miniopterus schreibersii* | Male | Spain | I (A. Daví) | 10/12/2012 | Few reads |
| M08013 | Blood | *Rhinolophus ferrumequinum* | Male | Morocco | II (Ghar-Knadel) | 25/06/2008 | Nearly full genome |
| M08017 | Blood | *Rhinolophus ferrumequinum* | Female | Morocco | II (Ghar-Knadel) | 25/06/2008 | Nearly full genome |
| M08051 | Blood | *Rhinolophus ferrumequinum* | Female | Morocco | II (Ghar-Knadel) | 25/06/2008 | Nearly full genome |
| M080113 | Blood | *Rhinolophus euryale* | Male | Morocco | III (Kef el Ghar) | 25/06/2008 | Few reads |
| M09005 | Blood | *Rhinolophus ferrumequinum* | Male | Morocco | IV (Ifri N' Caid) | 13/05/2009 | Nearly full genome |
| M09009 | Blood | *Rhinolophus ferrumequinum* | Male | Morocco | IV (Ifri N' Caid) | 13/05/2009 | Nearly full genome |
| A08011 | Blood | *Rhinolophus ferrumequinum* | Female | Algeria | V (Chrea) | 04/04/2008 | Nearly full genome |
| A09145 | Blood | *Rhinolophus ferrumequinum* | Male | Algeria | V (Chera) | 05/05/2009 | Nearly full genome |
| A09151 | Blood | *Rhinolophus ferrumequinum* | Male | Algeria | V (Chera) | 05/05/2009 | Nearly full genome |
| A09153 | Blood | *Rhinolophus ferrumequinum* | Male | Algeria | V (Chera) | 05/05/2009 | Nearly full genome |
| A09181 | Blood | *Rhinolophus ferrumequinum* | Male | Algeria | V (Chera) | 05/05/2009 | Nearly full genome |
| A09187 | Blood | *Rhinolophus ferrumequinum* | Female | Algeria | V (Chera) | 05/05/2009 | Few reads |
| A09193 | Blood | *Rhinolophus ferrumequinum* | Male | Algeria | V (Chera) | 05/05/2009 | Nearly full genome |
| A09197 | Blood | *Rhinolophus ferrumequinum* | Male | Algeria | V (Chera) | 05/05/2009 | Nearly full genome |
| A08065 | Blood | *Rhinolophus euryale* | Female | Algeria | VI (Aokas) | 06/04/2008 | Nearly full genome |
| A09061 | Blood | *Rhinolophus ferrumequinum* | Female | Algeria | VI (Aokas) | 02/05/2009 | Nearly full genome |
| A09097 | Blood | *Rhinolophus ferrumequinum* | Male | Algeria | VI (Aokas) | 03/05/2009 | Nearly full genome |

**Table S9**: Pairwise nucleotide comparison of the 16 bat rhabdovirus genomes expressed in nucleotide identity (%), after ORF concatenation for each of them.

| **Pairwise nucleotide identity (%)** | | | | | | | | | | | | | | | | |
| --- | --- | --- | --- | --- | --- | --- | --- | --- | --- | --- | --- | --- | --- | --- | --- | --- |
| **Concatenated ORFs** | **A09181** | **A09197** | **A09145** | **A09193** | **A08065** | **A09097** | **A09061** | **A09151** | **A09153** | **A08011** | **M09005** | **M09009** | **2012096** | **M08017** | **M08051** | **M08013** |
| **A09181** | 100 |  |  |  |  |  |  |  |  |  |  |  |  |  |  |  |
| **A09197** | 99.9 | 100 |  |  |  |  |  |  |  |  |  |  |  |  |  |  |
| **A09145** | 99.9 | 99.9 | 100 |  |  |  |  |  |  |  |  |  |  |  |  |  |
| **A09193** | 99.9 | 99.9 | 99.9 | 100 |  |  |  |  |  |  |  |  |  |  |  |  |
| **A08065** | 99.2 | 99.2 | 99.2 | 99.2 | 100 |  |  |  |  |  |  |  |  |  |  |  |
| **A09097** | 99.8 | 99.8 | 99.8 | 99.8 | 99.4 | 100 |  |  |  |  |  |  |  |  |  |  |
| **A09061** | 99.7 | 99.7 | 99.7 | 99.7 | 99.3 | 99.9 | 100 |  |  |  |  |  |  |  |  |  |
| **A09151** | 99 | 99 | 99 | 99 | 98.4 | 99 | 99 | 100 |  |  |  |  |  |  |  |  |
| **A09153** | 99 | 99 | 99 | 99 | 98.4 | 99 | 99 | 99.9 | 100 |  |  |  |  |  |  |  |
| **A08011** | 98.6 | 98.6 | 98.6 | 98.6 | 98.8 | 98.7 | 98.6 | 99.5 | 99.5 | 100 |  |  |  |  |  |  |
| **M09005** | 97.3 | 97.3 | 97.3 | 97.3 | 97.8 | 97.3 | 97.3 | 97.5 | 97.5 | 97.9 | 100 |  |  |  |  |  |
| **M09009** | 97.9 | 97.9 | 97.9 | 97.9 | 97.4 | 98 | 97.9 | 98.1 | 98.1 | 97.8 | 99.3 | 100 |  |  |  |  |
| **2012096** | 97.8 | 97.8 | 97.8 | 97.9 | 97.3 | 97.9 | 97.8 | 98 | 98 | 97.6 | 99.2 | 99.8 | 100 |  |  |  |
| **M08017** | 97.5 | 97.5 | 97.5 | 97.5 | 97.9 | 97.5 | 97.5 | 97.6 | 97.6 | 98 | 99 | 98.7 | 98.5 | 100 |  |  |
| **M08051** | 97.9 | 97.9 | 97.9 | 98 | 97.4 | 98 | 97.9 | 98.1 | 98.1 | 97.7 | 98.5 | 99.1 | 99 | 99.5 | 100 |  |
| **M08013** | 97.9 | 97.9 | 97.9 | 98 | 97.4 | 98 | 97.9 | 98.1 | 98.1 | 97.7 | 98.5 | 99.1 | 99 | 99.5 | 99.9 | 100 |

**Table S10.** Amino acid identities (%) of the P, G and L proteins between the 16 isolates of Mediterranean bat virus (MBV). Identities were calculated as pairwise deletion using MEGA7.0.

|  | **Pairwise amino acid identity (%)** | | | | | | | | | | | | | | | |
| --- | --- | --- | --- | --- | --- | --- | --- | --- | --- | --- | --- | --- | --- | --- | --- | --- |
|  | **2012096** | **M09005** | **M09009** | **M08013** | **M08017** | **M08051** | **A08011** | **A08065** | **A09061** | **A09097** | **A09145** | **A09151** | **A09153** | **A09181** | **A09193** | **A09197** |
| **P protein** |  |  |  |  |  |  |  |  |  |  |  |  |  |  |  |  |
| 2012096 | 100 |  |  |  |  |  |  |  |  |  |  |  |  |  |  |  |
| M09005 | 100 | 100 |  |  |  |  |  |  |  |  |  |  |  |  |  |  |
| M09009 | 100 | 100 | 100 |  |  |  |  |  |  |  |  |  |  |  |  |  |
| M08013 | 99.1 | 99.1 | 99.1 | 100 |  |  |  |  |  |  |  |  |  |  |  |  |
| M08017 | 99.1 | 99.1 | 99.1 | 100 | 100 |  |  |  |  |  |  |  |  |  |  |  |
| M08051 | 99.1 | 99.1 | 99.1 | 100 | 100 | 100 |  |  |  |  |  |  |  |  |  |  |
| A08011 | 98.6 | 98.6 | 98.6 | 98.6 | 98.6 | 98.6 | 100 |  |  |  |  |  |  |  |  |  |
| A08065 | 99.1 | 99.1 | 99.1 | 99.1 | 99.1 | 99.1 | 99.5 | 100 |  |  |  |  |  |  |  |  |
| A09061 | 99.1 | 99.1 | 99.1 | 99.1 | 99.1 | 99.1 | 99.5 | 100 | 100 |  |  |  |  |  |  |  |
| A09097 | 99.1 | 99.1 | 99.1 | 99.1 | 99.1 | 99.1 | 99.5 | 100 | 100 | 100 |  |  |  |  |  |  |
| A09145 | 99.1 | 99.1 | 99.1 | 99.1 | 99.1 | 99.1 | 99.5 | 100 | 100 | 100 | 100 |  |  |  |  |  |
| A09151 | 98.6 | 98.6 | 98.6 | 98.6 | 98.6 | 98.6 | 100 | 99.5 | 99.5 | 99.5 | 99.5 | 100 |  |  |  |  |
| A09153 | 98.6 | 98.6 | 98.6 | 98.6 | 98.6 | 98.6 | 100 | 99.5 | 99.5 | 99.5 | 99.5 | 100 | 100 |  |  |  |
| A09181 | 99.1 | 99.1 | 99.1 | 99.1 | 99.1 | 99.1 | 99.5 | 100 | 100 | 100 | 100 | 99.5 | 99.5 | 100 |  |  |
| A09193 | 99.1 | 99.1 | 99.1 | 99.1 | 99.1 | 99.1 | 99.5 | 100 | 100 | 100 | 100 | 99.5 | 99.5 | 100 | 100 |  |
| A09197 | 99.1 | 99.1 | 99.1 | 99.1 | 99.1 | 99.1 | 99.5 | 100 | 100 | 100 | 100 | 99.5 | 99.5 | 100 | 100 | 100 |
| **G protein** |  |  |  |  |  |  |  |  |  |  |  |  |  |  |  |  |
| 2012096 | 100 |  |  |  |  |  |  |  |  |  |  |  |  |  |  |  |
| M09005 | 99.6 | 100 |  |  |  |  |  |  |  |  |  |  |  |  |  |  |
| M09009 | 99.8 | 99.8 | 100 |  |  |  |  |  |  |  |  |  |  |  |  |  |
| M08013 | 99.4 | 99.4 | 99.6 | 100 |  |  |  |  |  |  |  |  |  |  |  |  |
| M08017 | 99.4 | 99.4 | 99.6 | 100 | 100 |  |  |  |  |  |  |  |  |  |  |  |
| M08051 | 99.4 | 99.4 | 99.6 | 100 | 100 | 100 |  |  |  |  |  |  |  |  |  |  |
| A08011 | 99 | 99 | 99.2 | 98.8 | 98.8 | 98.8 | 100 |  |  |  |  |  |  |  |  |  |
| A08065 | 98.8 | 98.8 | 99 | 98.6 | 98.6 | 98.6 | 99.8 | 100 |  |  |  |  |  |  |  |  |
| A09061 | 98.8 | 98.8 | 99 | 98.6 | 98.6 | 98.6 | 99.8 | 100 | 100 |  |  |  |  |  |  |  |
| A09097 | 98.8 | 98.8 | 99 | 98.6 | 98.6 | 98.6 | 99.8 | 100 | 100 | 100 |  |  |  |  |  |  |
| A09145 | 98.8 | 98.8 | 99 | 98.6 | 98.6 | 98.6 | 99.8 | 100 | 100 | 100 | 100 |  |  |  |  |  |
| A09151 | 99 | 99 | 99.2 | 98.8 | 98.8 | 98.8 | 100 | 99.8 | 99.8 | 99.8 | 99.8 | 100 |  |  |  |  |
| A09153 | 99 | 99 | 99.2 | 98.8 | 98.8 | 98.8 | 100 | 99.8 | 99.8 | 99.8 | 99.8 | 100 | 100 |  |  |  |
| A09181 | 98.8 | 98.8 | 99 | 98.6 | 98.6 | 98.6 | 99.8 | 100 | 100 | 100 | 100 | 99.8 | 99.8 | 100 |  |  |
| A09193 | 98.8 | 98.8 | 99 | 98.6 | 98.6 | 98.6 | 99.8 | 100 | 100 | 100 | 100 | 99.8 | 99.8 | 100 | 100 |  |
| A09197 | 98.8 | 98.8 | 99 | 98.6 | 98.6 | 98.6 | 99.8 | 100 | 100 | 100 | 100 | 99.8 | 99.8 | 100 | 100 | 100 |
| **L protein** |  |  |  |  |  |  |  |  |  |  |  |  |  |  |  |  |
| 2012096 | 100 |  |  |  |  |  |  |  |  |  |  |  |  |  |  |  |
| M09005 | 99.7 | 100 |  |  |  |  |  |  |  |  |  |  |  |  |  |  |
| M09009 | 99.8 | 99.9 | 100 |  |  |  |  |  |  |  |  |  |  |  |  |  |
| M08013 | 99.7 | 99.8 | 99.9 | 100 |  |  |  |  |  |  |  |  |  |  |  |  |
| M08017 | 99.7 | 99.8 | 99.9 | 100 | 100 |  |  |  |  |  |  |  |  |  |  |  |
| M08051 | 99.7 | 99.8 | 99.9 | 100 | 100 | 100 |  |  |  |  |  |  |  |  |  |  |
| A08011 | 99 | 99.1 | 99.2 | 99.2 | 99.2 | 99.2 | 100 |  |  |  |  |  |  |  |  |  |
| A08065 | 99.1 | 99.2 | 99.3 | 99.3 | 99.3 | 99.3 | 99.5 | 100 |  |  |  |  |  |  |  |  |
| A09061 | 99.1 | 99.2 | 99.3 | 99.4 | 99.4 | 99.4 | 99.5 | 99.9 | 100 |  |  |  |  |  |  |  |
| A09097 | 99.1 | 99.2 | 99.3 | 99.4 | 99.4 | 99.4 | 99.5 | 99.9 | 100 | 100 |  |  |  |  |  |  |
| A09145 | 99.1 | 99.2 | 99.3 | 99.3 | 99.3 | 99.3 | 99.5 | 99.9 | 99.9 | 99.9 | 100 |  |  |  |  |  |
| A09151 | 99 | 99 | 99.1 | 99.2 | 99.2 | 99.2 | 99.8 | 99.5 | 99.6 | 99.6 | 99.5 | 100 |  |  |  |  |
| A09153 | 99 | 99 | 99.1 | 99.2 | 99.2 | 99.2 | 99.8 | 99.5 | 99.6 | 99.6 | 99.5 | 100 | 100 |  |  |  |
| A09181 | 99.1 | 99.2 | 99.3 | 99.3 | 99.3 | 99.3 | 99.5 | 99.9 | 99.9 | 99.9 | 100 | 99.5 | 99.5 | 100 |  |  |
| A09193 | 99.1 | 99.2 | 99.3 | 99.3 | 99.3 | 99.3 | 99.5 | 99.9 | 99.9 | 99.9 | 100 | 99.5 | 99.5 | 100 | 100 |  |
| A09197 | 99 | 99.1 | 99.2 | 99.3 | 99.3 | 99.3 | 99.4 | 99.8 | 99.9 | 99.9 | 99.9 | 99.5 | 99.5 | 99.9 | 99.9 | 100 |

**Table S11.** Amino acid identities (%) of the 5 proteins (N, P, M, G and L) of Mediterranean bat virus (isolate 2012096) between other species among the genus *Vesiculovirus*. Identities were calculated as pairwise deletion using MEGA7.0.

|  | **Pairwise amino acid identity (%)** | | | | | | | | | | | | | | | | | | | |
| --- | --- | --- | --- | --- | --- | --- | --- | --- | --- | --- | --- | --- | --- | --- | --- | --- | --- | --- | --- | --- |
|  | **MBV (2012096)** | **YSBV (1017)** | **QZBV (1127)** | **JHBV** | **ABV** | **YBV** | **CARV** | **VSIV** | **VSAV** | **COCV** | **MORV** | **RADIV** | **MSPV** | **VSNJV** | **MARAV** | **JURV** | **PIRV** | **ISFV** | **CHNV** | **PERV** |
| **N Protein** |  |  |  |  |  |  |  |  |  |  |  |  |  |  |  |  |  |  |  |  |
| MBV_2012096 | 100 |  |  |  |  |  |  |  |  |  |  |  |  |  |  |  |  |  |  |  |
| YSBV_1017 | 75.5 | 100 |  |  |  |  |  |  |  |  |  |  |  |  |  |  |  |  |  |  |
| QZBV_1127 | 74.6 | 89.8 | 100 |  |  |  |  |  |  |  |  |  |  |  |  |  |  |  |  |  |
| JHBV | 74.6 | 89.5 | 99.7 | 100 |  |  |  |  |  |  |  |  |  |  |  |  |  |  |  |  |
| ABV | 53.5 | 54.2 | 54.7 | 54.4 | 100 |  |  |  |  |  |  |  |  |  |  |  |  |  |  |  |
| YBV | 47.8 | 46.7 | 47.4 | 47.4 | 48.5 | 100 |  |  |  |  |  |  |  |  |  |  |  |  |  |  |
| CARV | 46.9 | 45.5 | 45.5 | 45.5 | 46.4 | 53.3 | 100 |  |  |  |  |  |  |  |  |  |  |  |  |  |
| VSIV | 46.4 | 46.9 | 46.9 | 46.7 | 45.5 | 52.9 | 75.8 | 100 |  |  |  |  |  |  |  |  |  |  |  |  |
| VSAV | 46.2 | 45.5 | 45.5 | 45.3 | 44.6 | 53.3 | 74.8 | 84.8 | 100 |  |  |  |  |  |  |  |  |  |  |  |
| COCV | 46 | 45.5 | 45.7 | 45.5 | 45.3 | 51.7 | 73.9 | 83.6 | 85.3 | 100 |  |  |  |  |  |  |  |  |  |  |
| MORV | 46 | 47.1 | 45.5 | 45.5 | 46.2 | 52.2 | 74.8 | 90.5 | 83.4 | 83.4 | 100 |  |  |  |  |  |  |  |  |  |
| RADIV | 46 | 45.5 | 43.2 | 43.4 | 47.4 | 74.5 | 53.1 | 54.7 | 53.8 | 51.7 | 53.6 | 100 |  |  |  |  |  |  |  |  |
| MSPV | 45.5 | 46.9 | 45.8 | 45.8 | 46.5 | 51.9 | 55.2 | 55.2 | 54.3 | 55 | 53.8 | 54.2 | 100 |  |  |  |  |  |  |  |
| VSNJV | 45.5 | 47.4 | 46.2 | 46.2 | 46.9 | 52.9 | 72.5 | 69.1 | 69.4 | 70.1 | 68.7 | 54.5 | 55.5 | 100 |  |  |  |  |  |  |
| MARAV | 45.3 | 45 | 45 | 44.8 | 45.7 | 52.2 | 73.4 | 90 | 82.4 | 86.7 | 88.8 | 53.3 | 55.7 | 69.6 | 100 |  |  |  |  |  |
| JURV | 44.1 | 46.2 | 45.7 | 45.7 | 48.5 | 54 | 52.4 | 54.1 | 54.8 | 52.4 | 54.3 | 55.3 | 70.3 | 54.1 | 53.4 | 100 |  |  |  |  |
| PIRV | 43.7 | 46 | 43.9 | 43.9 | 48.6 | 51.2 | 49.5 | 52.5 | 51.4 | 50.9 | 52.1 | 53.9 | 61.5 | 51.4 | 51.6 | 62 | 100 |  |  |  |
| ISFV | 43.3 | 45.9 | 43.7 | 43.7 | 47.7 | 53.6 | 51.1 | 51.8 | 52.1 | 51.1 | 51.8 | 54.3 | 65.2 | 51.8 | 51.1 | 66.2 | 60.9 | 100 |  |  |
| CHNV | 42.8 | 42.8 | 42.6 | 42.6 | 47 | 48.9 | 50.2 | 50.4 | 48.1 | 48.1 | 50.4 | 50.9 | 58.1 | 50.2 | 50 | 61.4 | 55 | 58.3 | 100 |  |
| PERV | 38.3 | 38 | 36.6 | 36.6 | 39.6 | 42.5 | 45 | 45.3 | 45 | 45.3 | 45.3 | 44 | 52.8 | 46.2 | 45.7 | 49.8 | 51.6 | 47 | 43.7 | 100 |
| **P protein** |  |  |  |  |  |  |  |  |  |  |  |  |  |  |  |  |  |  |  |  |
| MBV_2012096 | 100 |  |  |  |  |  |  |  |  |  |  |  |  |  |  |  |  |  |  |  |
| YSBV_1017 | 21.1 | 100 |  |  |  |  |  |  |  |  |  |  |  |  |  |  |  |  |  |  |
| QZBV_1127 | 26.9 | 46 | 100 |  |  |  |  |  |  |  |  |  |  |  |  |  |  |  |  |  |
| JHBV | 27.8 | 48.8 | 86.6 | 100 |  |  |  |  |  |  |  |  |  |  |  |  |  |  |  |  |
| ABV | 11.6 | 9.7 | 9.4 | 9.8 | 100 |  |  |  |  |  |  |  |  |  |  |  |  |  |  |  |
| YBV | 12.3 | 14.8 | 13.6 | 12.9 | 9.4 | 100 |  |  |  |  |  |  |  |  |  |  |  |  |  |  |
| CARV | 12.7 | 9.5 | 10.7 | 11.7 | 14.6 | 11.9 | 100 |  |  |  |  |  |  |  |  |  |  |  |  |  |
| VSIV | 12.9 | 8.6 | 11.2 | 11.2 | 14.5 | 16.1 | 46.5 | 100 |  |  |  |  |  |  |  |  |  |  |  |  |
| VSAV | 10.1 | 8.1 | 11 | 11.3 | 15 | 15.8 | 44.4 | 53.5 | 100 |  |  |  |  |  |  |  |  |  |  |  |
| COCV | 10.4 | 10.4 | 10.9 | 11.3 | 17.5 | 16.4 | 44.6 | 58.2 | 60.2 | 100 |  |  |  |  |  |  |  |  |  |  |
| MORV | 13.5 | 8.9 | 11.8 | 11.8 | 16.2 | 18.8 | 45.4 | 67.7 | 53.7 | 61.8 | 100 |  |  |  |  |  |  |  |  |  |
| RADIV | 12.8 | 9.8 | 11.8 | 10 | 11 | 36.3 | 15.1 | 15.3 | 14.7 | 16.4 | 14.5 | 100 |  |  |  |  |  |  |  |  |
| MSPV | 10.3 | 11 | 10.1 | 10.1 | 15.6 | 12.4 | 16.4 | 17.4 | 16.4 | 18.1 | 17.6 | 15 | 100 |  |  |  |  |  |  |  |
| VSNJV | 10.1 | 8.4 | 8.8 | 9.8 | 14 | 11.2 | 37.8 | 29.8 | 32.4 | 33.2 | 30.8 | 12.8 | 15.1 | 100 |  |  |  |  |  |  |
| MARAV | 12.6 | 9.7 | 10.9 | 10.9 | 16 | 18.1 | 46.8 | 62.1 | 58.2 | 62 | 64.4 | 15.3 | 15.7 | 33 | 100 |  |  |  |  |  |
| JURV | 8.9 | 8.7 | 10 | 10.6 | 14.1 | 13.6 | 17.1 | 14.6 | 13 | 14.9 | 13.9 | 14 | 20.8 | 17.4 | 13.3 | 100 |  |  |  |  |
| PIRV | 11.4 | 9.6 | 10.2 | 11.1 | 14.1 | 12.5 | 22.3 | 16.3 | 13.8 | 14.5 | 16 | 11.3 | 19.2 | 17.5 | 17.2 | 30.2 | 100 |  |  |  |
| ISFV | 7.3 | 7.7 | 10.4 | 10.7 | 13.6 | 13.6 | 15.5 | 14.9 | 15.9 | 14.9 | 13.9 | 11.2 | 17.5 | 15.5 | 13.6 | 28.7 | 26.7 | 100 |  |  |
| CHNV | 11 | 9.7 | 9.8 | 9.1 | 14.5 | 13.9 | 18.4 | 18.3 | 19.1 | 19.1 | 16.2 | 13.5 | 22.8 | 14.4 | 16.4 | 22.9 | 26.6 | 23.8 | 100 |  |
| PERV | 8.2 | 7.3 | 12.4 | 10.4 | 14.6 | 11.2 | 15.5 | 15.8 | 17.2 | 16.5 | 15.8 | 13.7 | 22.9 | 18.8 | 14.9 | 31.2 | 26.6 | 27.6 | 25 | 100 |
| **M protein** |  |  |  |  |  |  |  |  |  |  |  |  |  |  |  |  |  |  |  |  |
| MBV_2012096 | 100 |  |  |  |  |  |  |  |  |  |  |  |  |  |  |  |  |  |  |  |
| YSBV_1017 | 61.5 | 100 |  |  |  |  |  |  |  |  |  |  |  |  |  |  |  |  |  |  |
| QZBV_1127 | 56.7 | 73.5 | 100 |  |  |  |  |  |  |  |  |  |  |  |  |  |  |  |  |  |
| JHBV | 56.2 | 74.5 | 95.1 | 100 |  |  |  |  |  |  |  |  |  |  |  |  |  |  |  |  |
| ABV | 40.4 | 40.9 | 39.5 | 40 | 100 |  |  |  |  |  |  |  |  |  |  |  |  |  |  |  |
| YBV | 18.9 | 19.7 | 19.3 | 20.1 | 23.8 | 100 |  |  |  |  |  |  |  |  |  |  |  |  |  |  |
| CARV | 19.2 | 18.8 | 18.4 | 17.9 | 19.2 | 17.7 | 100 |  |  |  |  |  |  |  |  |  |  |  |  |  |
| VSIV | 18.7 | 17.9 | 18.7 | 19.6 | 19.2 | 20 | 55.6 | 100 |  |  |  |  |  |  |  |  |  |  |  |  |
| VSAV | 18.3 | 17.9 | 19.2 | 19.6 | 22.7 | 21.2 | 55.6 | 74.6 | 100 |  |  |  |  |  |  |  |  |  |  |  |
| COCV | 19.6 | 18.3 | 20 | 20.5 | 21.3 | 18.8 | 57.3 | 75.1 | 81.2 | 100 |  |  |  |  |  |  |  |  |  |  |
| MORV | 18.3 | 17.4 | 17.9 | 18.3 | 20 | 19.2 | 60 | 82.9 | 77.2 | 79 | 100 |  |  |  |  |  |  |  |  |  |
| RADIV | 20.9 | 19.2 | 19.2 | 19.6 | 26.3 | 54.1 | 20.3 | 19.5 | 21.5 | 20.3 | 21.1 | 100 |  |  |  |  |  |  |  |  |
| MSPV | 22 | 22.8 | 23.2 | 24.1 | 25.8 | 27.7 | 23.8 | 27.2 | 26.8 | 25.9 | 26.3 | 29.4 | 100 |  |  |  |  |  |  |  |
| VSNJV | 18.3 | 15.2 | 17.9 | 18.7 | 18.7 | 20 | 54.5 | 62.1 | 60.8 | 60.8 | 62.6 | 20.2 | 24.5 | 100 |  |  |  |  |  |  |
| MARAV | 18.3 | 17 | 19.2 | 20 | 20.5 | 20 | 56.9 | 79.9 | 77.2 | 79.9 | 82.9 | 22.3 | 26.3 | 63.9 | 100 |  |  |  |  |  |
| JURV | 21.9 | 21.3 | 20.5 | 20.5 | 27 | 29.5 | 27.4 | 26.6 | 26.6 | 26.1 | 28.3 | 28.3 | 41.6 | 29.4 | 28.3 | 100 |  |  |  |  |
| PIRV | 21 | 23.4 | 20.8 | 21.2 | 29.7 | 30.7 | 21.3 | 25.9 | 26.7 | 26.3 | 26.3 | 29.1 | 39.9 | 24.5 | 25.5 | 46.5 | 100 |  |  |  |
| ISFV | 22.1 | 20.7 | 22 | 22.9 | 28.1 | 32.3 | 23.8 | 26.3 | 26.8 | 27.2 | 25.9 | 32.9 | 43.8 | 27.1 | 27.2 | 53.7 | 47.4 | 100 |  |  |
| CHNV | 20.7 | 20.6 | 21.1 | 20.6 | 27.5 | 29.5 | 23.7 | 25 | 26.2 | 24.5 | 27.1 | 31.2 | 36.5 | 23.6 | 26.2 | 47.5 | 41.8 | 50.2 | 100 |  |
| PERV | 17.9 | 19 | 19.9 | 19.9 | 22.8 | 27.8 | 23.2 | 23.2 | 26.1 | 23.6 | 24.4 | 27.3 | 38.3 | 26 | 26.1 | 46 | 48.3 | 46.8 | 40.1 | 100 |
| **G protein** |  |  |  |  |  |  |  |  |  |  |  |  |  |  |  |  |  |  |  |  |
| MBV_2012096 | 100 |  |  |  |  |  |  |  |  |  |  |  |  |  |  |  |  |  |  |  |
| YSBV_1017 | 62.4 | 100 |  |  |  |  |  |  |  |  |  |  |  |  |  |  |  |  |  |  |
| QZBV_1127 | 59.3 | 71.9 | 100 |  |  |  |  |  |  |  |  |  |  |  |  |  |  |  |  |  |
| JHBV | 59.5 | 71.7 | 84.8 | 100 |  |  |  |  |  |  |  |  |  |  |  |  |  |  |  |  |
| ABV | 31.2 | 29.6 | 30.9 | 30.9 | 100 |  |  |  |  |  |  |  |  |  |  |  |  |  |  |  |
| YBV | 24.2 | 24.3 | 24.1 | 24.3 | 23.9 | 100 |  |  |  |  |  |  |  |  |  |  |  |  |  |  |
| CARV | 25.9 | 27.5 | 28.3 | 28.7 | 26.5 | 35.2 | 100 |  |  |  |  |  |  |  |  |  |  |  |  |  |
| VSIV | 27.5 | 29 | 28.6 | 28.6 | 23.4 | 34.1 | 54.1 | 100 |  |  |  |  |  |  |  |  |  |  |  |  |
| VSAV | 27.5 | 27.5 | 28.4 | 28.2 | 22.2 | 33.5 | 53.7 | 62.3 | 100 |  |  |  |  |  |  |  |  |  |  |  |
| COCV | 26.9 | 28 | 28.7 | 28.5 | 22.6 | 32.6 | 54.1 | 71.2 | 66.9 | 100 |  |  |  |  |  |  |  |  |  |  |
| MORV | 27.6 | 27.8 | 27.6 | 28 | 24.1 | 35.1 | 55.6 | 84.4 | 63.9 | 71.5 | 100 |  |  |  |  |  |  |  |  |  |
| RADIV | 24.9 | 24.9 | 24.2 | 24 | 23.9 | 67 | 34.9 | 34.3 | 33.8 | 34.1 | 35.5 | 100 |  |  |  |  |  |  |  |  |
| MSPV | 26.7 | 27.4 | 25.3 | 25.5 | 26.1 | 40.2 | 36.9 | 35.6 | 37.1 | 36.3 | 37 | 38.1 | 100 |  |  |  |  |  |  |  |
| VSNJV | 26.4 | 27.5 | 27.5 | 27.3 | 23.8 | 31.4 | 50.5 | 49.2 | 47.9 | 47.2 | 48.8 | 32 | 36.9 | 100 |  |  |  |  |  |  |
| MARAV | 27.7 | 28.1 | 28.6 | 28.4 | 23.4 | 33.7 | 54.6 | 77.3 | 63.9 | 73.8 | 77.4 | 35.2 | 36.1 | 49.7 | 100 |  |  |  |  |  |
| JURV | 26.3 | 27.8 | 27 | 26.4 | 25.2 | 41.8 | 38.5 | 36.4 | 36.8 | 38.7 | 37.3 | 40.1 | 48.6 | 36.7 | 37.1 | 100 |  |  |  |  |
| PIRV | 27 | 27.5 | 26 | 25.6 | 26.1 | 44.2 | 40.5 | 37.6 | 36.8 | 36.8 | 37.4 | 42.9 | 47.7 | 37 | 38.2 | 48.8 | 100 |  |  |  |
| ISFV | 27 | 27.9 | 26.4 | 26.4 | 24.4 | 40.1 | 38.4 | 38.4 | 36 | 40 | 39.6 | 39.2 | 47.9 | 37.1 | 37.5 | 53.5 | 50 | 100 |  |  |
| CHNV | 27.4 | 26.7 | 26.1 | 26.1 | 23.7 | 41.9 | 41.1 | 38.3 | 39.2 | 39.6 | 39.8 | 41.5 | 48.3 | 37 | 39.8 | 52.4 | 51.5 | 54.1 | 100 |  |
| PERV | 27.7 | 28 | 26.5 | 26.7 | 24.8 | 43.3 | 38.7 | 37.7 | 38.9 | 37.3 | 38.2 | 41.6 | 49 | 38.2 | 37.9 | 51 | 56.5 | 49.1 | 50.9 | 100 |
| **L protein** |  |  |  |  |  |  |  |  |  |  |  |  |  |  |  |  |  |  |  |  |
| MBV_2012096 | 100 |  |  |  |  |  |  |  |  |  |  |  |  |  |  |  |  |  |  |  |
| YSBV_1017 | 67 | 100 |  |  |  |  |  |  |  |  |  |  |  |  |  |  |  |  |  |  |
| QZBV_1127 | 66 | 75.5 | 100 |  |  |  |  |  |  |  |  |  |  |  |  |  |  |  |  |  |
| JHBV | 65.9 | 76 | 91.3 | 100 |  |  |  |  |  |  |  |  |  |  |  |  |  |  |  |  |
| ABV | 56.8 | 57.5 | 57.6 | 57 | 100 |  |  |  |  |  |  |  |  |  |  |  |  |  |  |  |
| YBV | 53.9 | 53.2 | 52.7 | 53 | 54.8 | 100 |  |  |  |  |  |  |  |  |  |  |  |  |  |  |
| CARV | 53.1 | 53 | 52.8 | 52.5 | 54 | 57.8 | 100 |  |  |  |  |  |  |  |  |  |  |  |  |  |
| VSIV | 52.5 | 51.9 | 52.2 | 52.1 | 53.4 | 56.7 | 69.7 | 100 |  |  |  |  |  |  |  |  |  |  |  |  |
| VSAV | 53.5 | 53.2 | 52.4 | 52.1 | 54 | 57.4 | 69.7 | 75.5 | 100 |  |  |  |  |  |  |  |  |  |  |  |
| COCV | 52.6 | 53.3 | 52.5 | 52.5 | 53.2 | 57.7 | 68.9 | 76.4 | 78.2 | 100 |  |  |  |  |  |  |  |  |  |  |
| MORV | 52.5 | 52.8 | 53.3 | 52.8 | 53.1 | 57.3 | 69 | 80.3 | 75.3 | 77.2 | 100 |  |  |  |  |  |  |  |  |  |
| RADIV | 54.7 | 54.5 | 53.5 | 53.2 | 55.9 | 72.3 | 57.5 | 56.8 | 57.4 | 57.8 | 56.7 | 100 |  |  |  |  |  |  |  |  |
| MSPV | 53.9 | 54.2 | 55.1 | 54.5 | 56.7 | 59.3 | 59.2 | 58.9 | 59 | 59.1 | 58.4 | 61 | 100 |  |  |  |  |  |  |  |
| VSNJV | 52.5 | 52.4 | 52.4 | 52.2 | 53.8 | 56.6 | 69.7 | 65.8 | 67.3 | 65.6 | 65.9 | 57.4 | 57.9 | 100 |  |  |  |  |  |  |
| MARAV | 52.6 | 52.3 | 52 | 51.9 | 52.6 | 57.2 | 68.5 | 78 | 77.1 | 78.9 | 78.7 | 56.8 | 58.4 | 65.7 | 100 |  |  |  |  |  |
| JURV | 52.9 | 53.4 | 53.2 | 53.2 | 56.9 | 58.1 | 58.4 | 58.1 | 57.9 | 57.9 | 57.2 | 60.1 | 67.4 | 57.5 | 57.6 | 100 |  |  |  |  |
| PIRV | 53 | 52.8 | 53.7 | 53.2 | 56.9 | 59.2 | 58.7 | 57.3 | 57.6 | 57 | 57.3 | 60.1 | 67.6 | 57.9 | 56.9 | 66.5 | 100 |  |  |  |
| ISFV | 52.9 | 54.8 | 53.9 | 53.9 | 55.6 | 58.6 | 58.8 | 58.2 | 58.3 | 58.4 | 58.3 | 61 | 68.6 | 57.5 | 58.5 | 66.1 | 66.4 | 100 |  |  |
| CHNV | 53.3 | 54.3 | 54.5 | 54 | 55.3 | 59.8 | 58.6 | 58.8 | 58.2 | 57.8 | 58 | 61.6 | 67.4 | 57.6 | 58.5 | 66.5 | 66.7 | 68.5 | 100 |  |
| PERV | 52.4 | 53.7 | 53.5 | 53.5 | 54.8 | 58.3 | 58.7 | 57.1 | 57.3 | 57.8 | 58.1 | 59.6 | 66.7 | 57.1 | 57.3 | 64.5 | 66.9 | 66.5 | 66.3 | 100 |
